# PhyloMix: Enhancing microbiome-trait association prediction through phylogeny-mixing augmentation

**DOI:** 10.1101/2024.08.26.609661

**Authors:** Yifan Jiang, Disen Liao, Qiyun Zhu, Yang Young Lu

## Abstract

**Motivation:** Understanding the associations between traits and microbial composition is a fundamental objective in microbiome research. Recently, researchers have turned to machine learning (ML) models to achieve this goal with promising results. However, the effectiveness of advanced ML models is often limited by the unique characteristics of microbiome data, which are typically high-dimensional, compositional, and imbalanced. These characteristics can hinder the models’ ability to fully explore the relationships among taxa in predictive analyses. To address this challenge, data augmentation has become crucial. It involves generating synthetic samples with artificial labels based on existing data and incorporating these samples into the training set to improve ML model performance.

**Results:** Here we propose PhyloMix, a novel data augmentation method specifically designed for microbiome data to enhance predictive analyses. PhyloMix leverages the phylogenetic relationships among microbiome taxa as an informative prior to guide the generation of synthetic microbial samples. Leveraging phylogeny, PhyloMix creates new samples by removing a subtree from one sample and combining it with the corresponding subtree from another sample. Notably, PhyloMix is designed to address the compositional nature of microbiome data, effectively handling both raw counts and relative abundances. This approach introduces sufficient diversity into the augmented samples, leading to improved predictive performance. We empirically evaluated PhyloMix on six real microbiome datasets across five commonly used ML models. PhyloMix significantly outperforms distinct baseline methods including sample-mixing-based data augmentation techniques like vanilla mixup and compositional cutmix, as well as the phylogeny-based method TADA. We also demonstrated the wide applicability of PhyloMix in both supervised learning and contrastive representation learning.

**Availability:** The Apache licensed source code is available at (https://github.com/batmen-lab/phylomix).

## 1 Introduction

The human microbiome represents the intricate communities of microorganisms residing in and on our bodies, with bacteria alone possessing 100 times more unique genes than the human genome (Qin et al, 2010). These microbiomes play a crucial role in human physiology, significantly affecting both health and disease (Sekirov et al, 2010). For example, the microbiome influences a wide range of diseases, including diabetes (Qin et al, 2012), obesity (Turnbaugh et al, 2006), inflammatory bowel disease (Mills et al, 2022), and even neurodegenerative conditions such as Parkinson’s disease (Dutta et al, 2019) and Alzheimer’s disease (Vogt et al, 2017). Exploring the association between the microbiome and host traits will offer insights into the mechanisms by which the microbiome influences human health and disease, paving the way for the development of innovative therapeutic strategies.

To investigate microbiome-trait associations, researchers have employed machine learning (ML) models to uncover complex patterns within microbial samples under various conditions and thereby facilitate microbiome-based disease detection (McCoubrey et al, 2021). Disease prediction from microbial samples is typically framed as a supervised learning task. In this approach, an ML model is trained on microbial samples–often characterized by the abundance of selected microbial taxa–to predict a host trait of interest, which may be binary (*e*.*g*., disease status) or continuous (*e*.*g*., age). Extensive studies have shown that ML models hold significant potential for accurately classifying clinically relevant traits across various diseases using microbial sample profiles (Wu et al, 2021; Medina et al, 2022). These ML models range from classic approaches like logistic regression and support vector machines to more advanced techniques like random forests and deep neural networks.

Despite increasing interest, the effectiveness of advanced ML models has been constrained by the unique characteristics of microbiome data, limiting their ability to fully explore the relationships among taxa in predictive analyses. Specifically, microbiome features, which represent the abundance of hundreds to thousands of taxa, are inherently high-dimensional. This high dimensionality presents analytical challenges, as the relatively small number of samples can result in overfitting during training, thus limiting the model’s ability to generalize to unseen samples or even leading to conflicting research conclusions (Knights et al, 2011; Reiman et al, 2020). Furthermore, microbiome data is inherently compositional because the profiling instrument imposes an arbitrary total on the profiled taxa abundances (Gloor et al, 2017). This can lead to issues when compositional data is analyzed using non-compositional methods. Lastly, the host traits of interest are often unevenly distributed, with training data either over-representing diseased states and lacking sufficient healthy samples, or vice versa, resulting in a distribution that does not accurately reflect the general population. Thus, new methods are required to adapt machine learning models to complex data challenges, including low sample sizes, high feature dimensionality, compositionality, and unbalanced trait distribution.

Ideally, ML models could be better trained for predictive analyses with more microbial samples, but in reality, collecting these samples is laborious and costly. From a practitioner’s perspective, a more cost-effective alternative is to computationally augment the data, an approach that has been increasingly studied by the ML community in recent years (Mumuni and Mumuni, 2022). The core concept of data augmentation is to generate synthetic samples with artificial labels based on existing data, and incorporate these into the training data to enhance ML model performance (Fig. 1**a**). For example, one of the pioneering methods, SMOTE (Chawla et al, 2002), generates synthetic samples by oversampling minority classes using k-nearest intra-class neighbors. More recently, mixup (Zhang et al, 2018) and its variants (Cao et al, 2022) create synthetic samples by linearly combining two samples and their corresponding labels, which are drawn randomly from the training data. However, most previous works are general-purpose and not tailored for microbiome data, overlooking domain-specific knowledge unique to the microbiome. Numerous approaches have been developed to augment microbiome data, with generative methods such as PhylaGAN (Sharma et al, 2024), MB-GAN (Rong et al, 2021), and TADA (Sayyari et al, 2019) standing out as notable advancements. These techniques aim to capture the data’s underlying distributional characteristics. While GAN-based methods are powerful, they often face challenges in effectively training on small-scale datasets due to their high sensitivity to data quantity and quality. To our knowledge, the only exception is TADA (Sayyari et al, 2019), but this algorithm is limited to mimicking sample-wise variations and fails to fully explore the potential of advanced data augmentation techniques (See detailed comparison in Sec. 4).

**Fig. 1:**
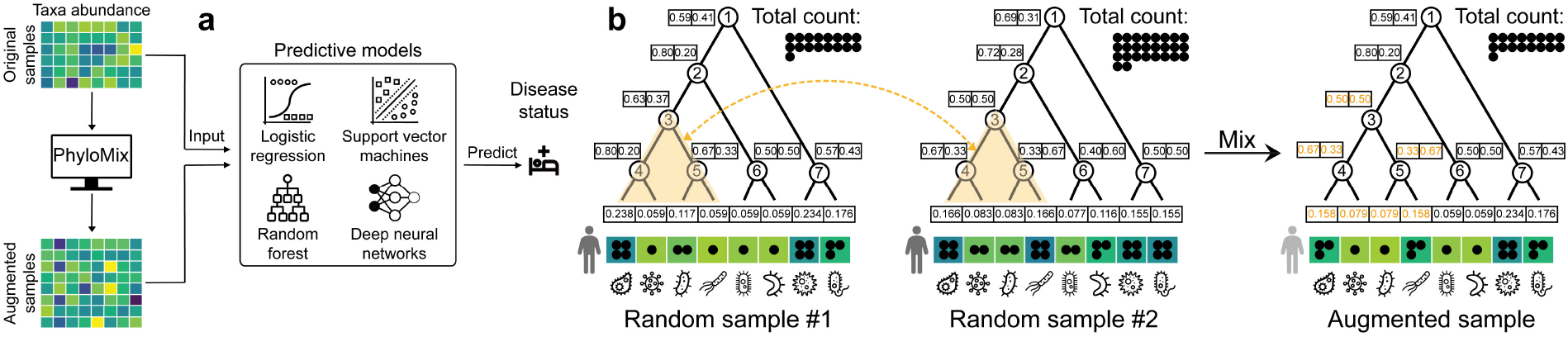
Overview of PhyloMix. (**a**) PhyloMix generates synthetic samples with artificial labels based on existing data and incorporates them into the training data to enhance the performance of general ML models. (**b**) PhyloMix leverages phylogeny to generate synthetic samples by removing a subtree from one sample and recombining it with the counterpart from another sample. The involved subtree, along with the affected probabilistic phylogenetic profile, is highlighted in orange.

In this study, we introduce a novel data augmentation method specifically designed for microbiome data to enhance predictive analyses, called PhyloMix (<u>Phylo</u>geny-<u>Mix</u>ing augmentation). The key ideas behind PhyloMix is to leverage the phylogenetic relationships among microbiome taxa, using them as an informative prior to guide the generation of synthetic microbial samples. Here, phylogeny serves as a tool to organize and understand microbial taxa by encoding their evolutionary relationships. Leveraging phylogeny, PhyloMix generates new samples by removing a subtree from one sample and recombining the counterpart from another sample, rather than interpolating between two whole microbial samples. This approach considers that variations among microbiome taxa are constrained by their evolutionary relationships as encoded by the phylogeny, thereby producing more realistic synthetic samples. Importantly, PhyloMix encodes phylogenetic relationships using a novel phylogenetic probabilistic profiling to address the compositional nature of microbiome data, allowing it to effectively handle both raw counts and relative abundances.

We have evaluated PhyloMix for supervised learning tasks using three simulated and six publicly available microbiome datasets, each with varying sample sizes and feature dimensionality. We also bench-marked the evaluation using five commonly used ML models, ranging from classic logistic regression and support vector machines to more advanced methods like random forests and deep neural networks. Overall, the augmentation generated by PhyloMix preserves or notably enhances predictive performance, regardless of the dataset complexity or the sophistication of the ML models used. In addition to supervised learning, PhyloMix enables representation learning by using randomly augmented training samples to define a self-supervised optimization objective, pulling random augmentations of the same sample closer together while pushing those generated from different samples further apart (Chen et al, 2020). We observe that PhyloMix outperforms previous representation learning approaches by identifying more distinguishable representations of samples with different host trait labels.

## 2 Approach

### 2.1 Problem setup

Consider a supervised learning task involving a training data of *n* microbial samples, represented as a data matrix. The rows of the data matrix correspond to the set of samples, denoted as 𝒳= {*x*_1_, *x*_2_, …, *x*_*n*_ }where *x*_*i*_ ∈ ℝ^*p*^ represents the *p*-dimensional features of the *i*-th sample. The columns of the data matrix represent the set of features corresponding to the abundance of microbial taxa, denoted as 𝒯 = {*t*_1_, *t*_2_, …, *t*_*p*_} where *t*_*j*_ ∈ ℝ^*n*^ represents the abundance of the *j*-th taxon observed across all samples. The abundance of microbial taxa can be measured either by count or by normalized probability. The training data also includes trait labels, denoted as 𝒴 = {*y*_1_, *y*_2_, …}, *y*_*n*_ . Each sample *x*_*i*_ is paired with a corresponding trait label *y*_*i*_ ∈ ℝ, which is binary(*e*.*g*., disease status) or continuous (*e*.*g*., age).

Additionally, we have phylogenetic relationships among microbial taxa that encode their evolutionary dependencies. The phylogeny classifies taxa through a series of splits, representing estimated events where two lineages diverge from a common ancestor to form distinct species (Washburne et al, 2018). Two taxa with close phylogenetic relationships are likely to have similar functional roles, exhibiting more similar abundance variations compared to two randomly selected taxa. PhyloMix uses an inferred binary phylogenetic tree (See Sec. 3.2), denoted as 𝒫, with the tree’s leaf nodes labeled by the set of taxa 𝒯. The tree’s internal nodes are indexed from 1 to *p*−1 in a top-down fashion, denoted as 𝒱 = {*v*_1_, *v*_2_, …, *v*_*p−*1_}.

Given the samples 𝒳, the corresponding trait labels 𝒳, as well as the phylogenetic tree 𝒫, our goal is to generate synthetic samples 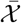 with corresponding synthetic trait labels 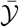, where 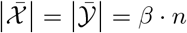 and *β >* 0 is the augmentation ratio specified by the user. Empowered by the augmented samples 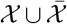 with labels 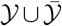, a more effective ML model *f* : ℝ^*p*^ ↦ ℝ should be learned (Fig. 1**a**), mapping a sample *𝒳* ∈ ℝ^*p*^ to the trait labels *𝒴* ∈ ℝ, which may no longer remains binary after the data augmentation.

### 2.2 Sample mixup

PhyloMix draws inspiration from mixup (Zhang et al, 2018), a widely-used data augmentation technique that creates synthetic samples by linearly combining two randomly drawn samples from the same dataset and their corresponding labels. Specifically, given two random drawn sample-label pairs (*x*_*i*_, *y*_*i*_) and (*x*_*j*_, *y*_*j*_), where *x*_*i*_, *x*_*j*_ ∈ 𝒳 denotes the input samples and *y*_*i*_, *y*_*j*_ ∈ 𝒴 are their corresponding trait labels, mixup a new synthetic sample 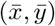 by:

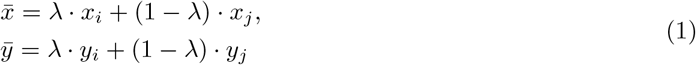

where *λ* ∈ [0, 1] is the mixing weight drawn from a beta distribution. It is worth noting that mixup is designed to be generic, treating the features corresponding to the abundance of microbial taxa equally. Considering that recent studies have leveraged the inherent correlation structure among taxa as an informative prior to improve disease prediction performance (Xiao et al, 2018; Albanese et al, 2015; Sharma et al, 2020; Reiman et al, 2020; Jiang et al, 2023), we hypothesize that a data augmentation technique specifically tailored for microbiome data by incorporating this information can further enhance predictive accuracy.

### 2.3 Constructing sample-specific phylogenetic profiles

PhyloMix aims to enhance mixup-based data augmentation by incorporating phylogenetic relationships among microbiome taxa to guide the generation of synthetic samples. We are aware of the mixup variation that uses constituency parsing trees to generate synthetic sentences in the natural language processing community (Zhang et al, 2022). However, this method cannot be directly applied to microbiome data due to its unique characteristics. Specifically, each microbial sample is represented by a set of features corresponding to the abundance of microbial taxa, measured either by count or normalized probability. In the case of count data, two samples may be profiled under different sequencing depths. Thus, a higher count measured for taxon A in sample 1 compared to taxon B in sample 2 may result either from taxon A being more highly expressed than B or from sample 1 having a much higher overall sequencing depth than sample 2. In the case of normalized probability, where counts across taxa are scaled to sum to 1, substituting the probability of taxon A in sample 1 with that of sample 2 may violate the sum-to-one constraint in sample 1, rendering the resultant features of the synthetic sample invalid.

To address this challenge, we construct a probabilistic phylogenetic profile for each sample, ensuring that it remains valid to remove features corresponding to an arbitrary subtree from one sample and recombine them with the counterpart from another sample (Fig. 1**b**). Specifically, the probabilistic phylogenetic profile of a sample *x* includes the following information: (1) the total abundance 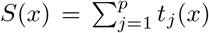, where *t*_*j*_(*x*) indicates the abundance of the *j*-th taxon in sample *x*, and (2) the phylogenetic probability (*l*_*v*_(*x*), *r*_*v*_(*x*)) for each internal node in the phylogenetic tree, where *l*_*v*_(*x*) and *r*_*v*_(*x*) indicate the relative abundance of the descendants of its left and right children, respectively, with *l*_*v*_(*x*) + *r*_*v*_(*x*) = 1:

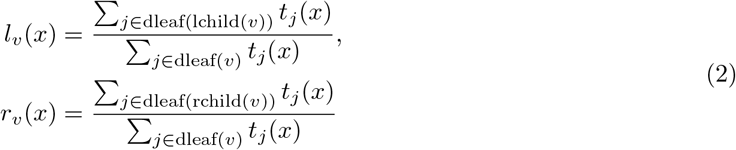

where lchild(*v*) ⊂ 𝒱 ∪ 𝒯 and rchild(*v*) ⊂ 𝒱 ∪ 𝒯 denote the left and right children of *v*, and dleaf(*v*) ⊂ 𝒯 denotes the descendent leaf nodes of *v*.

With the probabilistic phylogenetic profile of *x*, we can reconstruct the relative and absolute abundance for each microbiome taxon by traversing the phylogenetic tree in a top-down fashion. Specifically, for the *j*-th taxon, we trace the path from the root to the leaf node *t*_*j*_ and calculate its relative abundance by multiplying the phylogenetic probabilities of the internal nodes encountered along the path.

For example, as illustrated in Fig. 1**b**, the relative abundance of the first taxon in sample 1 is calculated as *p*_1_(*x*) = 0.59 × 0.80 × 0.63 × 0.80 ≈ 0.238. Similarly, the relative abundance of the last taxon in sample 1 is calculated as *p*_8_(*x*) = 0.41 × 0.43 ≈ 0.176. Given the total abundance of *x* across all taxa, the absolute abundance for the *j*-th taxon is simply the allocation of this total abundance to that taxon. For example, the absolute abundance of the first and last taxa in sample 1 is calculated as *t*_1_(*x*) = *p*_1_(*x*) × *S*(*x*) = 0.238 × 17 ≈ 4 and *t*_2_(*x*) = *p*_2_(*x*) × *S*(*x*) = 0.176 × 17 ≈ 3, respectively.

### 2.4 Generating synthetic samples

With probabilistic phylogenetic profiles enabling valid exchange of arbitrary subtree between samples, we can now leverage phylogenetic information to guide the generation of synthetic microbial samples. Concretely, for a given sample-label pair (*x*_*i*_, *y*_*i*_), we randomly sample another sample-label pair (*x*_*j*_, *y*_*j*_) from the training data. First, we construct probabilistic phylogenetic profiles for both samples, allowing us to sample subtrees for exchange. Next, we draw a mixing weight *λ* ∈ [0, 1] from a beta distribution Beta(*α, α*), where *α* ∈ (0, ∞) determines whether the synthetic sample combines evenly from the two original samples or is biased toward one of them. Intuitively, we want the portion of taxa exchanged to be reasonable. If the portion exchanged is too small or too large, the exchange may closely resemble the original samples and fail to introduce sufficient diversity into the augmented sample. Therefore, we use a Beta(2, 2) distribution in this study, ensuring that the mixing weights drawn are centered around 0.5 (See Sec. 4.3 for the robustness of different Beta distribution options).

To ensure that the taxa exchanged are constrained by the phylogenetic tree, we employ a two-phase sampling strategy. Specifically, We randomly draw an internal node *v* ∈ 𝒱 from the phylogenetic tree and exchange the phylogenetic profiles between (*l*_*v*_(*x*_*i*_), *r*_*v*_(*x*_*i*_)) and (*l*_*v*_(*x*_*j*_), *r*_*v*_(*x*_*j*_)) for all descendent internal nodes of *v*, provided they have not already been exchanged. We repeat this draw-and-exchange process the number of exchanged internal nodes approximately reaches (1 − *λ*) · *p*. Once we have the mixed probabilistic phylogenetic profiles, with *λ* · *p* internal nodes originally belonging to sample *x*_*i*_ and (1 − *λ*) · *p* to sample *x*_*j*_, we can reconstruct the relative and absolute abundance for each microbiome taxon based on the total abundance *S*(*x*_*i*_), as described in Sec. 2.3. We denote the synthetic sample with the generated absolute abundance values as 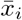, and create its label as 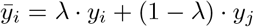. We repeat the process of generating synthetic samples until we have *β · n* new samples, where *β >* 0 is the augmentation ratio specified by the user. We use *β* = 3 in this study (See Sec. 4.3 for the robustness of different *β* options).

## 3 Materials and methods

### 3.1 Datasets

We conducted a comprehensive evaluation of PhyloMix using six publicly available microbiome datasets with varying sample sizes and feature dimensionality (Details in Tab. A.1). The AlzBiom dataset (Laske et al, 2022) investigated the connection between the gut microbiome and Alzheimer’s disease (AD). It includes 75 amyloid-positive AD samples and 100 cognitively healthy control samples from the AlzBiom study, with profiles containing *p* = 8, 350 taxa. The ASD dataset (Dan et al, 2020) examined the relationship between the gut microbiome and abnormal metabolic activity in individuals with Autism Spectrum Disorder (ASD). It consists of 30 typically developing (TD) and 30 constipated ASD (C-ASD) samples, with profiles containing *p* = 7, 287 taxa. The GD dataset (Zhu et al, 2021) investigated the link between the gut microbiome and Graves’ disease (GD). It includes 100 GD samples and 62 healthy control samples, with profiles containing *p* = 8, 487 taxa. The RUMC dataset (Boktor et al, 2023) is part of a cohort study exploring the relationship between the gut microbiome and the Parkinson’s disease (PD). The dataset includes 42 PD samples and 72 healthy control samples, with profiles containing *p* = 7, 256 taxa. The IBD dataset (Gonzalez et al, 2022) studied the relationship between the gut microbiome and two main subtypes of inflammatory bowel disease (IBD): Crohn’s disease (CD) and ulcerative colitis (UC). It includes 108 CD and 66 UC samples, with profiles containing *p* = 5, 287 taxa. The HMP2 dataset (Lloyd-Price et al, 2019), part of the Integrative Human Microbiome Project (iHMP) (, iHMP), also explored the relationship between the gut microbiome and two IBD subtypes: CD and UC. Compared to the IBD dataset, this dataset increases the sample size to *n* = 1, 158 (728 CD and 430 UC samples) and expands the taxa set to *p* = 10, 614.

Additionally, we created three simulated datasets using the microbiome data simulator MIDASim (He et al, 2024). We used a microbiome data simulator MIDASim (He et al, 2024) to generate simulated data by leveraging a template microbiome dataset while preserving its correlation structure to maintain similarity. We selected the IBD dataset as the template and adjusted the simulation settings to create three datasets with varying levels of difficulty in distinguishing between labels. Since MIDASim is not designed to generate simulated samples with labels, we selected a real dataset with positive and negative labels. We then simulated samples separately for each label (positive and negative) before combining them into a unified simulated dataset. (Refer to Sec. A.2 for the details of simulated dataset generation).

### 3.2 Phylogenetic tree

The abundance features in these datasets were profiled using Metagenomic Shotgun (MGS) sequencing, adhering to the standard operating procedures for shotgun metagenomics as implemented on the Qiita platform (Gonzalez et al, 2018). Specifically, the raw sequencing reads were classified using Woltka v0.1.4 (Zhu et al, 2022) against the Web of Life (WoL) v2 database (Zhu et al, 2019), which includes 15, 953 microbial genomes. Note that a small number of internal nodes in the WoL phylogeny have more than two lineages (*i*.*e*., polytomies). To address this, we resolved the issue by creating an arbitrary dichotomous structure among the affected nodes. (Refer to Sec. A.4 for the robustness of different polytomy resolution strategies). Furthermore, because each dataset only profiles a subset of taxa in the WoL phylogeny, we used a pruned phytogenetic tree having the dataset-specific taxa as leaf nodes only.

### 3.3 Evaluation tasks

We assessed the effectiveness of PhyloMix in two distinct settings: supervised learning and representation learning. The supervised learning setting evaluates PhyloMix’s ability to generate augmented samples that improve predictive performance for trait labels. To assess the predictive performance, we employed two metrics: the Area Under the Receiver Operating Characteristic Curve (AUROC) and the Area Under the Precision-Recall Curve (AUPRC). Both metrics are widely used to evaluate predictive performance, with AUPRC being particularly well-suited for imbalanced datasets (Davis and Goadrich, 2006).

The representation learning setting assesses PhyloMix’s capability to generate augmented samples that facilitate the extraction of meaningful high-level features from raw data (Bengio et al, 2013). PhyloMix adopts a contrastive learning-based representation learning approach. Specifically, PhyloMix uses the same encoder architecture as the deep autoencoder, mapping representations to a space where the contrastive loss is applied (Chen et al, 2020). The contrastive loss function is designed to bring the representations of positive pairs of samples closer together while pushing the representations of negative pairs further apart. Here, positive pairs are defined as two synthetic samples generated by applying random data augmentations to the same training sample pairs, whereas negative pairs are synthetic samples generated from different sample pairs. To evaluate representation learning performance, we utilized the learned representations as input for a downstream classifier to carry out predictive tasks.

### 3.4 Benchmark settings

We incorporated PhyloMix with five ML models with varying predictive capabilities: logistic regression (LR), support vector machine (SVM) with a linear kernel, random forest (RF), multi-layer perceptron (MLP), and MIOSTONE, a state-of-the-art deep learning model that encodes taxonomy (Jiang et al, 2023). We used the Scikit-learn implementation (Pedregosa et al, 2011) with default settings for the LR, RF, and SVM models. The MLP model was configured with a pyramid-shaped architecture featuring two hidden layers of size 256 and 128, respectively. For the MIOSTONE model, we employed the default implementation settings. (Refer to Sec. A.3 for the details of ML model settings and trainings). For all methods, we preprocessed microbiome features using centered log-ratio transformation (Aitchison, 1982) prior to data augmentation. (Refer to Sec. A.3 and Sec. A.6.2 for the robustness of using other types of data transformation such as relative abundance normalization).

It is important to note that while the trait labels for the training samples are binary, they become continuous for the generated synthetic samples. For deep learning models like MLP and MIOSTONE, training is based on cross-entropy loss, so the continuous labels will not affect the learning process. Continuous labels provide more robust predictions, similar to the role of label smoothing (Zhang et al, 2021). For other models like LR, SVM, and RF, which are designed for discrete labels, we instead use their regression counterparts–linear regression, support vector regression, and random forest regression– to train on the augmented data with continuous labels. We also utilized the Scikit-learn implementation (Pedregosa et al, 2011) with default settings for these regression models.

### 3.5 Baseline methods

We compared PhyloMix’s performance against several distinct baseline methods, each generating an equivalent amount of synthetic samples to train the ML models alongside the original training data. The vanilla mixup (Zhang et al, 2018), a classic data augmentation method, creates synthetic samples by linearly combining two samples along with their corresponding labels. The compositional cutmix (Gordon-Rodriguez et al, 2022) extends the vanilla mixup to accommodate compositional data. TADA (Sayyari et al, 2019) generates synthetic samples by explicitly modeling biological and sampling variations, guided by the evolutionary relationships among taxa. MB-GAN (Rong et al, 2021) employs generative adversarial networks to learn from microbial abundance data and generate simulated abundances that closely mimic the original data.

## 4 Results

### 4.1 PhyloMix enhances supervised learning

We evaluated data augmentation performance on three simulated and six real microbiome datasets with varying sample and microbial taxa sizes. These datasets cover a range of disease prediction tasks, including Autism Spectrum Disorder, Alzheimer’s disease, Graves’ disease, Parkinson’s disease, and inflammatory bowel disease. Thus, they provide a comprehensive evaluation for predictive models of the human gut microbiome.

We evaluated the predictive performance using 5-fold cross-validation, training individual models on each training split and testing them on the corresponding test splits. The predictions from all test splits were concatenated, and the AUROC and AUPRC scores were calculated on the full dataset. For robustness, we repeated this process 20 times using different random seeds and reported the mean performance along with the 95% confidence intervals. Each repetition involved both data splitting and ML model training.

Our analysis shows that the augmentation generated by PhyloMix preserves or notably enhances predictive performance (Fig. 2**a** and Sec. A.6.1), regardless of the dataset complexity or the sophistication of the ML models used. On average across all ML models, PhyloMix achieves a 4.4%, 6.4%, and 6.7% improvement in test AUPRC on three simulated datasets. In comparison, vanilla mixup, the second-best data augmentation method, yields only a 3.2%, 4.3%, and 5.7% improvement in test AUPRC on three simulated datasets. For real datasets, PhyloMix achieves a 2.6%, 8.7%, 2.4%, and 4.1% improvement in test AUPRC on the RUMC, AlzBiom, IBD, and ASD datasets, respectively. In comparison, vanilla mixup yields only a 1.8%, 5.6%, 1.3%, and 2.0% improvement in test AUPRC on the RUMC, AlzBiom, IBD, and ASD datasets, respectively. Importantly, PhyloMix rarely hurt model performance, with drops observed in only 2 out of 35 cases, and the average decrease is typically just 0.3%. In comparison, vanilla mixup shows decreased performance in 8 out of 35 cases, with an average reduction of 0.8%. In the remaining GD and HMP2 datasets, PhyloMix achieves only a modest improvement of 0.1% and 0.1% in test AUPRC, as these datasets already exhibit near-perfect predictive performance, regardless of data augmentation. To rigorously quantify how much better is PhyloMix compared to the vanilla mixup, we compared the performance between PhyloMix and the vanilla mixup using one-tailed two-sample t-tests to calculate p-values. PhyloMix demonstrates significantly better performance (p-value ≤ 0.05) in terms of test AUPRC on five out of eight datasets with the logistic regression model, seven out of eight datasets with the support vector machine model, four out of eight datasets with the random forest model, two out of eight datasets with the multi-layer perceptron model, and four out of eight datasets with the MIOSTONE model. In other words, PhyloMix performs at least as well as vanilla mixup and is significantly better in more than half of the scenarios. A similar conclusion can be drawn based on AUROC performance (Fig. A.4 and Fig. A.6).

**Fig. 2:**
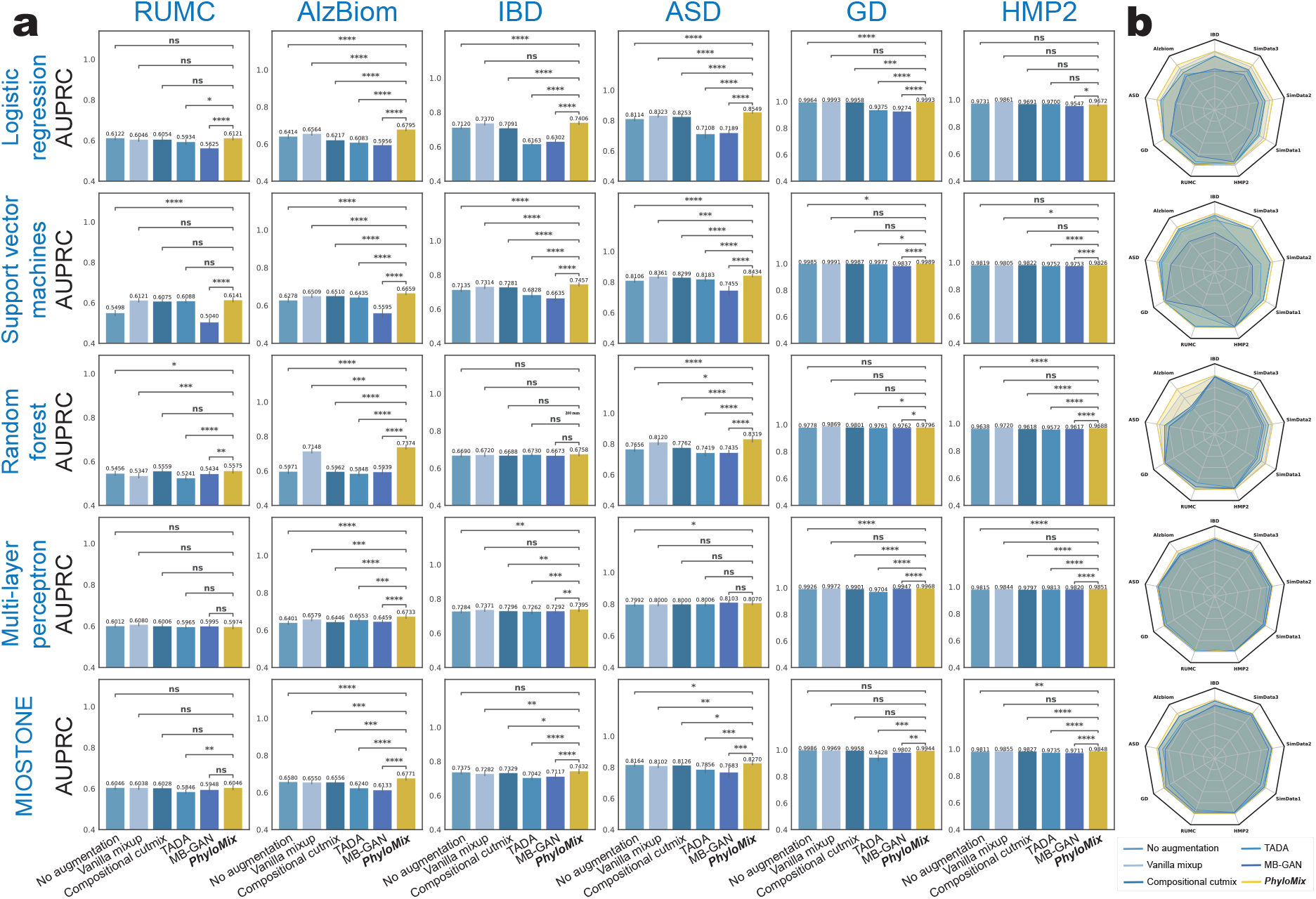
Data augmentation performance in the supervised learning setting. (**a**) The evaluation was performed on real microbiome datasets with varying microbial taxa sizes. PhyloMix is evaluated alongside five ML models with varying predictive capabilities and compared against four distinct baseline methods. The augmentation generated by PhyloMix preserves or notably enhances predictive performance, regardless of the dataset complexity or the sophistication of the ML models used. For scientific rigor, the performance comparison between PhyloMix and other baseline methods is quantified using one-tailed two-sample t-tests to calculate p-values: ∗ ∗ ∗ ∗ p-value ≤ 0.0001; ∗ ∗ ∗p-value ≤ 0.001; ∗ ∗ p-value ≤ 0.01; ∗ : p-value ≤ 0.05; ns : p-value > 0.05. (**b**) PhyloMix is compared to baseline methods using radar plots, with performance evaluated based on relative AUPRC, normalized against the bestperforming method across all methods. Each dot corresponds to the performance with or without data augmentation for a given method under a random seed.

Finally, in addition to the quantitative comparison, we conducted a qualitative evaluation of PhyloMix and other baseline methods (Fig. 2**b**. We directly visualize the improvement in predictive performance after applying each data augmentation method to generate synthetic training samples. The results demonstrate that PhyloMix consistently ranks among the top performers in both metrics across all datasets and various ML models. This is evident from the radar plots (Fig. 2**b**), where PhyloMix’s contours are always the largest and encompass those of other methods as subsets. We observed that TADA, which relies on modeling biological and sampling variations, fails to noticeably improve model performance, despite using the same phylogenetic information as PhyloMix. This underscores the importance of employing sample and label mixing for effective synthetic sample generation. Compositional cutmix and vanilla mixup can fail to enhance model performance when applied to LR and MIOSTONE models, highlighting their limitations in applicability. Importantly, PhyloMix consistently improves the performance of all ML models across different random seeds, regardless of the model’s complexity.

### 4.2 PhyloMix enhances representation learning

Having demonstrated improvements in predicting trait labels, we next evaluated PhyloMix’s capability to generate augmented samples that facilitate the extraction of meaningful high-level features from raw data. We compared PhyloMix’s performance against three alternative data augmentation methods: the vanilla mixup, the compositional cutmix, and TADA. In addition, since the UniFrac distance is a widely used metric for comparing microbial samples (Lozupone and Knight, 2005), we included an additional contrastive learning-based baseline, where positive and negative sample pairs were selected based on their UniFrac distance, with positive pairs having a smaller UniFrac distance compared to negative pairs. Furthermore, traditional representation learning typically employs a deep autoencoder to derive low-dimensional representations by minimizing reconstruction loss. For comparison, we included a deep autoencoder as a baseline method, where the encoder consists of three hidden layers with sizes of 512, 256, and 128, respectively. The decoder mirrors this architecture in reverse.

We assessed the quality of the learned representations using the following evaluation criteria. The learned representation from all methods is input into a logistic regression model to predict the trait label, using the same procedure as in the supervised learning setting. For robustness, we repeated this process 20 times using different random seeds and reported the mean performance along with the 95% confidence intervals. Each repetition involved data splitting, representation learning, and ML model training.

Our analysis reveals that PhyloMix’s contrastive model consistently outperforms both the deep autoencoder and the contrastive models that use alternative data augmentation methods in terms of learned representation quality (Fig. 3). On average, PhyloMix achieves a 9.8% and 9.8% improvement in test AUPRC and AUROC compared to using the representation learned by deep autoencoder, respectively. Importantly, PhyloMix enhances the quality of learned representations across all datasets, with the smallest improvement observed being 1.5% in test AUPRC and 3.0% in AUROC on the RUMC dataset. In comparison, compositional cutmix, the second-best data augmentation method in representation learning, shows only an 8.3% improvement in test AUPRC and an 8.2% improvement in AUROC over the deep autoencoder. Although UniFrac distance is widely used for comparing microbial samples based on phylogeny, PhyloMix proves to be more effective in leveraging phylogeny as an informative prior to enhance the quality of representation learning. The results demonstrate that PhyloMix consistently ranks among the top performers in both metrics across all datasets and various ML models.

**Fig. 3:**
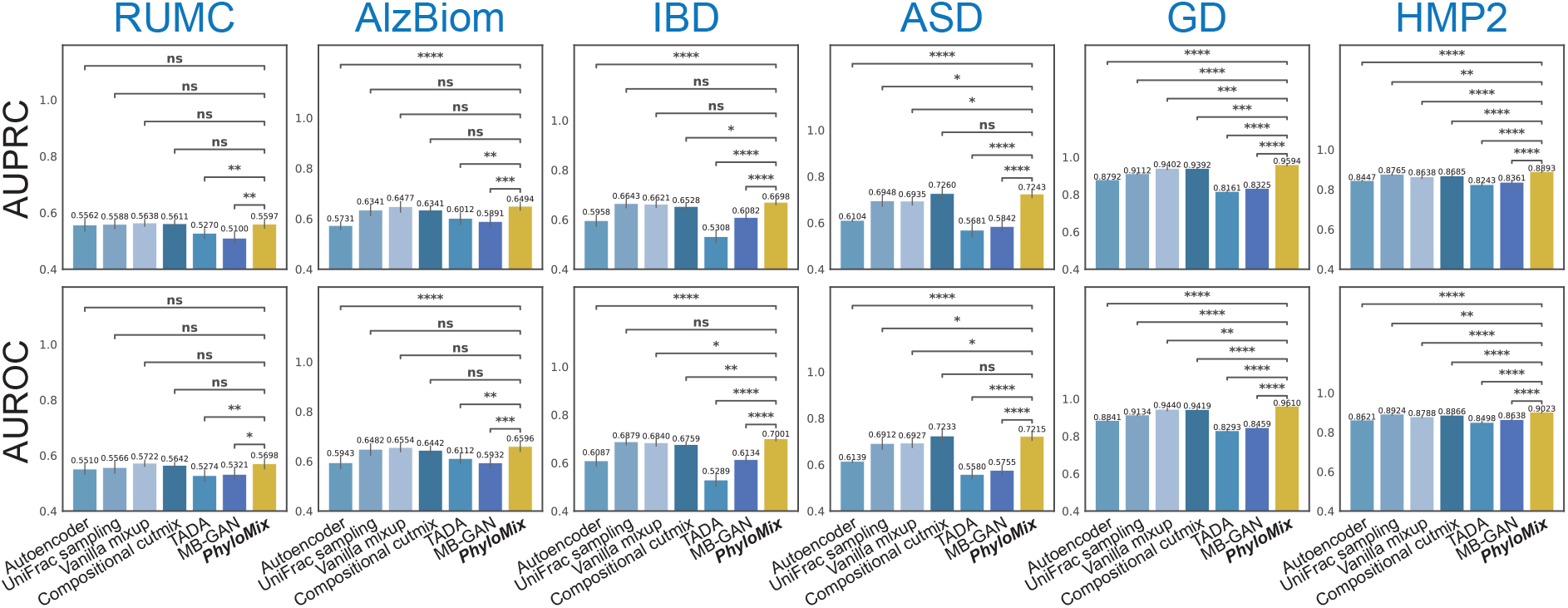
Data augmentation performance in the representation learning setting. The evaluation was conducted on six publicly available microbiome datasets and compared against six distinct baseline methods. The representations obtained through PhyloMix consistently outperform those generated by other baseline methods in predicting disease status, measured by both AUPRC and AUROC. The performance comparison between PhyloMix and other baseline methods is quantified using one-tailed two-sample t-tests to calculate p-values.

### 4.3 Dissecting the performance of PhyloMix

We first evaluated the computational cost of using PhyloMix in conjunction with five ML models with different levels of complexities. For a fair comparison, all models were trained and tested in the same environment: AMD EPYC 7302 16-Core Processor, NVIDIA RTX A6000 GPU, with 32GB DDR4 RAM. PhyloMix’s data augmentation involves two primary components: generating synthetic samples and the additional training time necessary to integrate these synthetic samples. We found that PhyloMix’s sample generation incurs only moderate computational overhead, measured in seconds (Fig. 4**a** and Fig. A.10). The additional training time required to incorporate these synthetic samples is marginal on the AlzBiom dataset, except for the HMP2 dataset, which has a larger sample size of *n* = 1, 158 and higher dimensionality with *p* = 10, 614 taxa. Note that when the sample size is large, data augmentation becomes less critical, as the dataset already exhibits near-perfect predictive performance, with or without augmentation. However, when the sample size is small, the computational time is negligible compared to the improvement in predictive accuracy gained through augmentation.

**Fig. 4:**
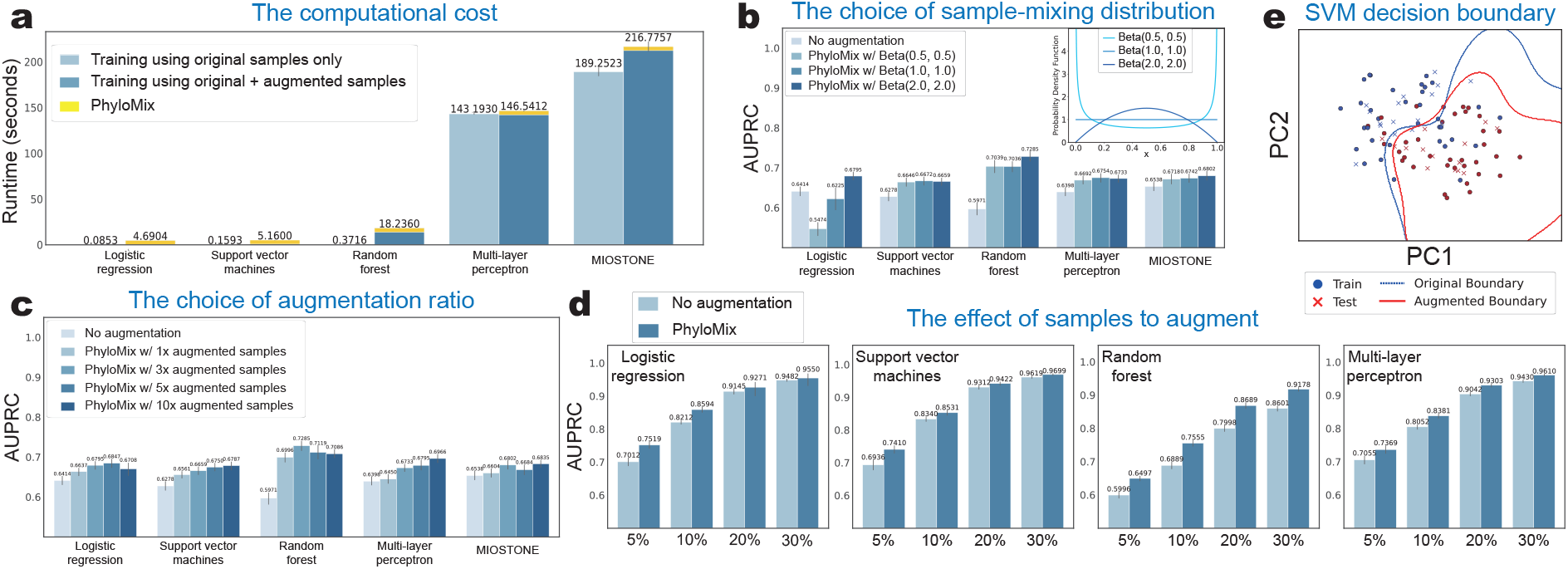
Dissecting the performance of PhyloMix through control studies. (**a**) The data augmentation by PhyloMix incurs only moderate computational overhead, consisting of two main components: the generation of synthetic samples and the additional training time required to incorporate these synthetic samples. (**b**) The choice of the Beta distribution is crucial in determining the sample mixing weights. In general, this choice does not negatively impact data augmentation performance. The only exception is the logistic regression model, where data augmentation transforms logistic regression into linear regression, leading to a performance downgrade if the Beta distribution is not chosen properly. (**c**) PhyloMix is robust to the number of augmented samples. Data augmentation consistently improves predictive performance. However, increasing the number of augmented samples inevitably leads to higher computational overhead. (**d**) PhyloMix demonstrates more significant improvements when the training data size is small. (**e**) PhyloMix smooths the decision boundary of the SVM model on the IBD dataset.

PhyloMix’s data augmentation relies on two factors: the choice of the sample-mixing distribution and the augmentation level. To evaluate the impact of each component on predictive performance, we conducted a control study using the AlzBiom dataset by modifying PhyloMix, altering its components with different variations Specifically, we considered two variants of PhyloMix: (1) Altering the Beta distributions that control sample mixing weights; (2) Adjusting the number of augmentation samples added. The results indicate that PhyloMix is robust to the choice of these two factors. In the first study (Fig. 4**b**), we compared three Beta distribution options, each representing a distinct sample mixing strategy. Specifically, Beta(0.5, 0.5) tends to bias the synthetic sample toward one of the original samples, Beta(2.0, 2.0) encourages a more even combination from both original samples, while Beta(1.0, 1.0) results in a more arbitrary combination. In general, we observed that the Beta distribution choice does not negatively impact data augmentation performance. The only exception is the logistic regression model, where data augmentation transforms logistic regression into linear regression, leading to a performance downgrade if the Beta distribution is not chosen properly so that the synthetic samples may closely resemble the original ones and fail to introduce sufficient diversity.

In the second study (Fig. 4**c**), we compared four different augmentation levels, ranging from adding the same number of synthetic samples as the original samples to adding up to ten times as many synthetic samples. In general, we observed that data augmentation consistently improves predictive performance. To achieve a significant performance boost, we expect the number of synthetic samples added to be at least three times the number of original samples. It’s important not to include excessive synthetic samples. On one hand, performance may diminish or even drop if too many synthetic samples are added; on the other hand, increasing the number of augmented samples leads to higher computational overhead. From a theoretical perspective, mixup-based data augmentation methods such as PhyloMix are theoretically analogous to certain regularization techniques, such as label smoothing and dropout (Carratino et al, 2022). (Refer to Sec. A.1 for more insights of data augmentation). From a regularization perspective, excessive regularization, such as generating an excessive number of synthetic samples, can have a detrimental effect by over-smoothing the labels, ultimately compromising prediction performance. Furthermore, we investigate the impact of varying sample sizes on PhyloMix’s predictive performance.

To address this question, we randomly selected subsets of training samples from the HMP2 dataset, with subset sizes ranging from 5%, 10%, 20%, and 30% of the total training samples, to train the ML models. For robustness, we repeated this process 20 times with different random seeds. For each taxa subset, we trained and evaluated PhyloMix using the same settings as for the full dataset. We found that while training PhyloMix with more samples generally leads to better predictive performance, the improvement provided by PhyloMix’s data augmentation is notably more significant across all ML models (Fig. 4**d**). It is important to note that data augmentation becomes more critical for performance when the sample size is small, highlighting the necessity of PhyloMix.

Finally, we examine the rationale behind PhyloMix’s data augmentation by visualizing its effect on smoothing the decision boundary of the SVM model. Specifically, we trained an SVM model on the training split of the IBD dataset, calculated the decision boundary, and examined how this boundary separates positive and negative samples in the test split (Fig. 4**e** and Fig. A.1). After data augmentation, the decision boundary shifts toward the mean of the output space, leading to a smoother and more generalized boundary. Note that the data augmentation is particularly effective at smoothing the decision boundary when the sample size is small. It is important to highlight that, beyond the empirical results, mixup-based data augmentation methods are grounded in strong theoretical principles (Carratino et al, 2022). (Refer to Sec. A.1 for more insights of data augmentation).

## 5 Discussion and conclusion

In this study, we propose PhyloMix, a novel data augmentation method specifically designed for microbiome data to enhance predictive analyses. At its core, PhyloMix leverages the phylogenetic relationships among microbiome taxa as an informative prior to guide the generation of synthetic microbial samples. Leveraging phylogeny, PhyloMix generates new samples by removing a subtree from one sample and recombining the counterpart from another sample, rather than interpolating between two whole microbial samples. The key novelties of PhyloMix are threefold: (1) We enable valid phylogeny exchange through the use of sample-specific probabilistic phylogenetic profiles, effectively handling both raw counts and relative abundances. (2) We demonstrate that phylogeny-based sample mixing achieves superior performance compared to relying solely on phylogeny or using sample mixing alone. (3) We showcase the wide applicability of PhyloMix in both supervised learning and contrastive representation learning.

We evaluated PhyloMix’s performance on simulated and real microbiome datasets, each with varying sample sizes and feature dimensionality. We benchmarked the evaluation using five commonly used ML models, ranging from classic logistic regression and support vector machines to more advanced methods like random forests and deep neural networks. Overall, the augmentation generated by PhyloMix preserves or notably enhances predictive performance, regardless of the dataset complexity or the sophistication of the ML models used. Furthermore, PhyloMix significantly outperforms distinct baseline methods in supervised learning and contrastive representation learning settings, including sample-mixing-based data augmentation techniques like vanilla mixup and compositional cutmix, as well as the phylogeny-based method TADA. Importantly, our comprehensive control studies demonstrate that PhyloMix is robust with respect to the choice of sample-mixing distribution and the level of augmentation.

Lastly, this study highlights several promising directions for future research. First, we observe that generating an excessive number of synthetic samples may have a detrimental effect and ultimately compromise prediction performance. An interesting future direction would be to determine the optimal number of synthetic samples for achieving the best prediction performance in a data-driven manner. Additionally, we emphasize how the synthetic data can quantitatively improve predictive performance. A potential future research direction is to explore how these synthetic data can contribute to generating meaningful biological insights and uncovering interesting biological phenomena. Finally, it is commonly believed that a better machine learning model is typically more interpretable (Adebayo et al, 2018). It is also observed that PhyloMix, along with other mixup-based data augmentation methods, can empirically improve the predictive performance of ML models. A promising direction for future research is to quantitatively assess the impact of data augmentation on the interpretability of machine learning models.

## Declarations

- The research is supported by the Canadian NSERC Discovery Grant RGPIN-03270-2023.
- The authors declare that they have no conflict of interest.
- There is no ethics approval and consent to participate involved in this study.
- There is no consent for publication involved in this study.
- Q.Z. provided the datasets, Y.J. and D.L implemented the code and did the analysis, D.L conducted the experiments, Y.L. and Y.J. prepared the figures. Y.L. wrote the main manuscript. All authors participated the discussion. All authors reviewed the manuscript.

## A Appendix

### A.1 The rationale of data augmentation

Our analysis demonstrates that across all datasets, mixup-based data augmentation methods–including the vanilla mixup (Zhang et al, 2018), the compositional cutmix (Gordon-Rodriguez et al, 2022), and PhyloMix–consistently outperform generative data augmentation approaches such as TADA (Sayyari et al, 2019) and MB-GAN (Rong et al, 2021). Beyond the empirical results, mixup-based data augmentation methods are theoretically analogous to certain regularization techniques, such as label smoothing and dropout (Carratino et al, 2022). This is due to their inherent ability to pull both inputs and outputs closer to their mean, which enhances model calibration and smooths the model’s Jacobian, ultimately improving generalization.

To illustrate the effect of data augmentation, we visualize how it smooths the decision boundary of the SVM model. Specifically, we trained an SVM model on the training split of the IBD dataset, calculated the decision boundary, and examined how this boundary separates positive and negative samples in the test split. As shown in Fig. A.1, after data augmentation, the decision boundary shifts toward the mean of the output space, leading to a smoother and more generalized boundary. Note that the data augmentation is particularly effective at smoothing the decision boundary when the sample size is small.

**Fig. A1:**
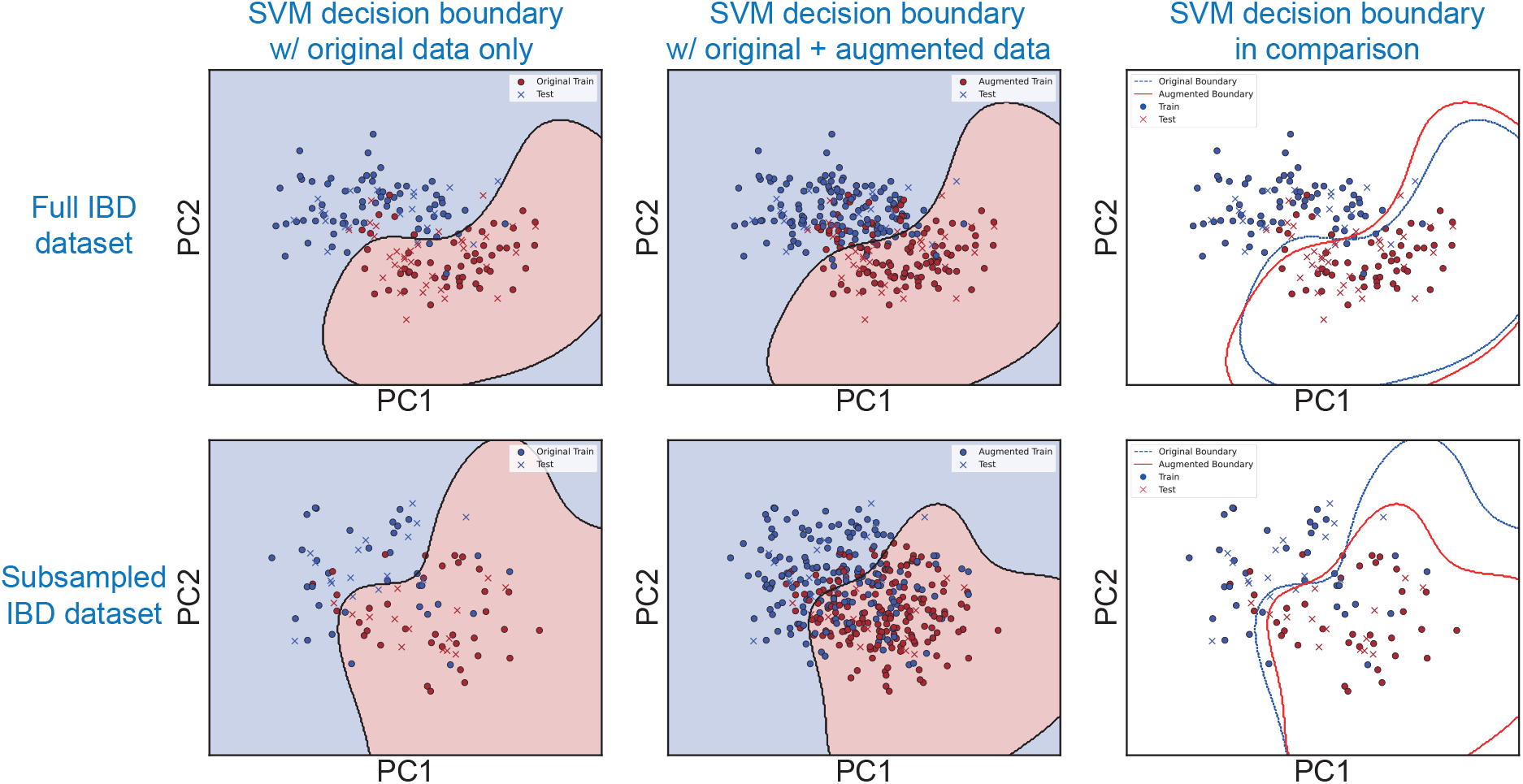
PhyloMix smooths the decision boundary of the SVM model. The evaluation was performed on the IBD dataset using both the full dataset and a subsampled version. Data augmentation is particularly effective at smoothing the decision boundary when the sample size is small.

### A.2 Dataset details

PhyloMix used six publicly available microbiome datasets with varying sample sizes and feature dimensionality, with details listed in Tab. A.1.

#### A.2.1 Simulated datasets

Additionally, we created three simulated datasets using the microbiome data simulator MIDASim (He et al, 2024). MIDASim generates simulated data by leveraging a template microbiome dataset and preserving its correlation structure to ensure similarity. Specifically, we employed the parametric mode of MIDASim, utilizing a generalized gamma distribution to model the relative abundances of microbiome data. This approach is tailored for simulation studies that involve modifying the log-mean relative abundance. Since MIDASim is not designed to generate simulated samples with labels, we selected a real dataset with positive and negative labels. We then simulated samples separately for each label (positive and negative) before combining them into a unified simulated dataset. The real dataset we chose is the IBD dataset (Gonzalez et al, 2022) studied the relationship between the gut microbiome and two main subtypes of inflammatory bowel disease (IBD): Crohn’s disease (CD) and ulcerative colitis (UC). It includes 108 CD and 66 UC samples, with profiles containing *p* = 5287 taxa.

**Table A1:** The details of the real datasets investigated by PhyloMix.

| Dataset | Sample size | Feature size | Data source | Notes |
| --- | --- | --- | --- | --- |
| AlzBiom | 175 samples (75 amyloid-positive and 100 healthy control) | 8,350 taxa | EBI-ENA ID: PRJEB47976 | The sequencing data is clean (QC'ed, host-filtered). |
| ASD | 60 samples (30 typically developing and 30 constipated ASD) | 7,287 taxa | EBI-ENA ID: PRJNA451479 | The sequencing data is raw (non-QC'ed). |
| GD | 162 samples (100 Graves' disease and 62 healthy control) | 8,487 taxa | EBI-ENA ID: PRJNA602729, PRJNA602731, PRJNA602732, PRJNA638403, PRJNA638404, PRJNA638405 | Most samples have a pair of FASTQ files. However, 4 samples (three.lst) have a third, unpaired FASTQ file that is very small, and it should be excluded from the analysis. 12 samples have only one FASTQ file, which appears to be single-end sequences. Two samples: GA61 (SRR12000211) and GA89 (SRR12005695) are missing from the metadata. Therefore they were dropped from the data. |
| RUMC | 114 samples (42 Parkinson's disease and 72 healthy control) | 7,256 taxa | Qiita ID: 12975 | 20 samples in BIOM are missing in metadata. These samples were dropped. |
| IBD | 174 samples (108 Crohn's disease and 66 ulcerative colitis) | 5,287 taxa | Qiita ID: 12675 | The dataset contains metagenomic sequencing data and associated metadata. More details can be found at: https://qiita.ucsd.edu/study/description/12675 |
| HMP2 | 1,158 samples (728 Crohn's disease and 430 ulcerative colitis) | 10,614 taxa | Qiita ID: 11484 | The dataset contains metagenomic sequencing data and associated metadata from the Human Microbiome Project. More details can be found at: https://hmpdacc.org/ihmp |

We simulated three distinct datasets from the IBD dataset. We estimated separate location parameters for each of the two labels from the IBD dataset, denoted as *µ*_+_ ∈ ℝ^5287^ and *µ*_*−*_ ∈ ℝ^5287^. After that, we varied the location parameters to represent different levels of difficulty in distinguishing between labels, as follows:

- Setting 1: (2 ∗ *µ*_+_) and (2 ∗*µ*_*−*_). (Alg. 1)
- Setting 2: (*µ*_+_ + 5) and (*µ*_*−*_ + 5). (Alg. 2)
- Setting 3: Randomly select 10% of taxa with non-zero abundance and increase their values by 10%. (Alg. 3)

Three settings introduce progressively greater challenges. Setting 1 involves minor deviations that preserve the presence-absence pattern but introduce moderate variability in relative abundances. In this setting, we doubled the mean location parameters of the gamma distribution for both positive and negative samples. This transformation preserves the presence-absence status of the taxa but shifts the location of the gamma distribution for each taxon. As a result, the relative abundance of taxa is altered due to the inherent randomness of the gamma distribution, though these changes remain relatively modest. Setting 2 significant shifts in distribution occur here, altering the relative abundance patterns substantially. In this setting, we shifted the mean location parameters by 5 units for both positive and negative samples. This adjustment ensures the presence of all taxa in the simulated dataset while significantly altering the underlying distribution. Consequently, the dataset exhibits greater deviations in relative abundance compared to the template, thereby increasing its complexity. In Setting 3, the original distribution structure is disrupted by directly modifying a subset of taxa, leading to pronounced deviations that may obscure the underlying data patterns. In this setting, the manipulation leads to substantial deviations from the original template. By altering the relative abundance of a subset of taxa, the overall structure of the dataset is disrupted, making this the most challenging scenario for model learning and inference. For each simulated dataset, we generated 100 positive samples and 100 negative samples.

##### Algorithm 1 Simulation Setting 1: Doubling the Location Parameter

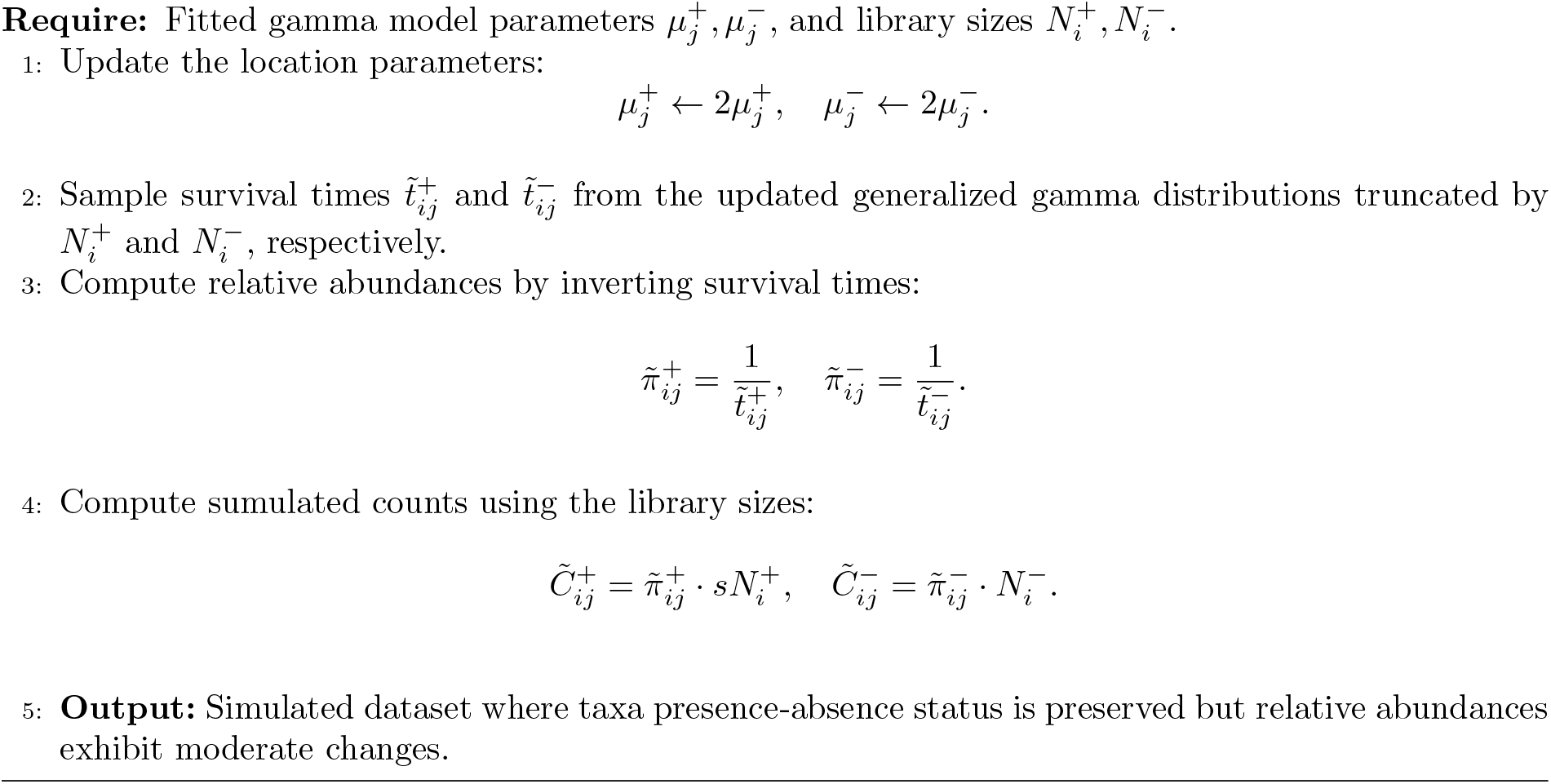

##### Algorithm 2 Simulation Setting 2: Shifting the Location Parameter

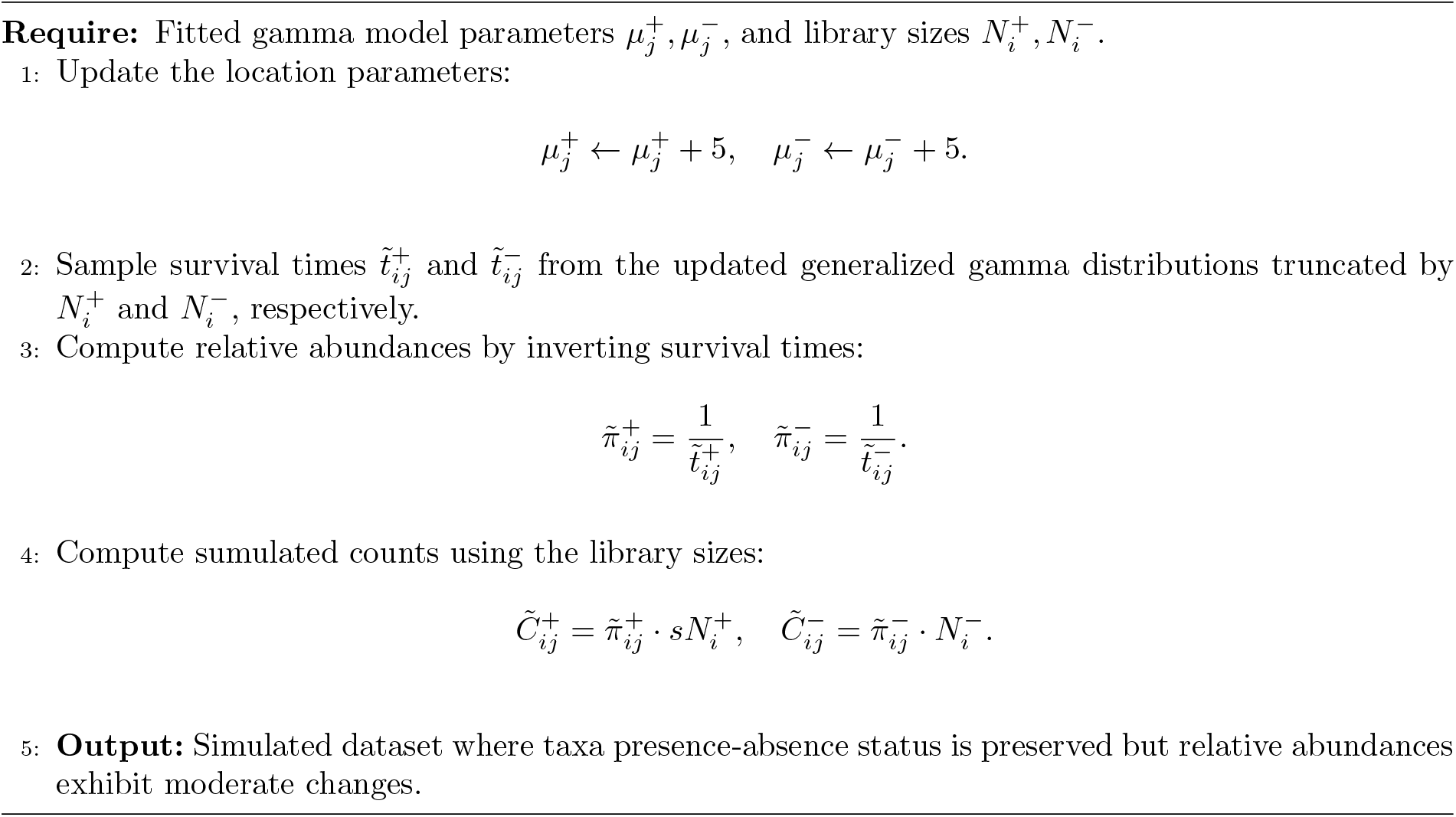

##### Algorithm 3 Simulation Setting 3: Modifying Relative Abundance

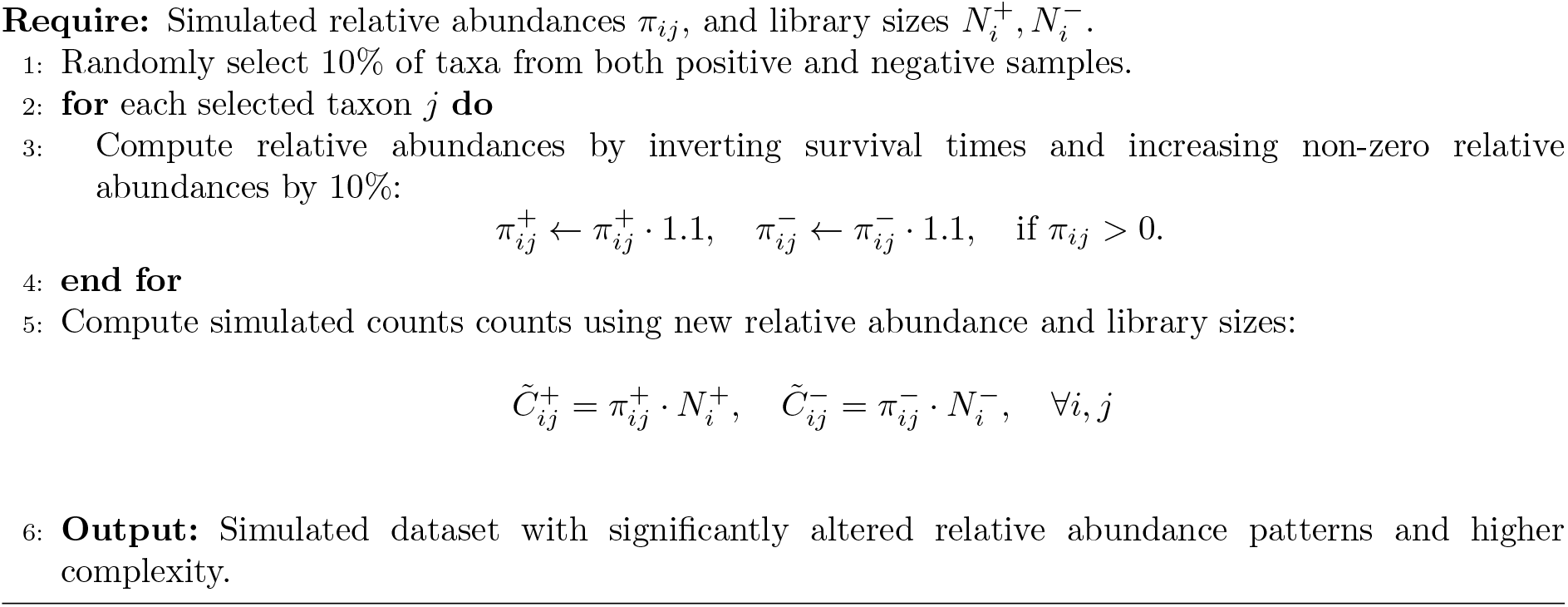

### A.3 Benchmark details

PhyloMix is used together with five ML models with varying predictive capabilities: logistic regression (LR), support vector machine (SVM) with a linear kernel, random forest (RF), multi-layer perceptron (MLP), and MIOSTONE, a state-of-the-art deep learning model that encodes taxonomy (Jiang et al, 2023). We used the Scikit-learn implementation (Pedregosa et al, 2011) with default settings for the LR, RF, and SVM models. Specifically, the logistic regression is trained using L2 regularization with a coefficient of 1.0. The random forest classifier is trained with 100 trees, without a maximum tree depth constraint, and a minimum of 2 samples required to split an internal node. The support vector classifier is trained with a linear kernel, using L2 regularization with a coefficient of 1.0. The MLP model was configured with a pyramid-shaped architecture featuring two hidden layers of size 256 and 128, respectively. For the MIOSTONE model, we employed the default implementation settings. We trained both the MLP and MIOSTONE models for 200 epochs with a batch size of 512 to ensure convergence. During training, we used the AdamW optimizer with a learning rate of 0.001 and applied a cosine annealing scheduler. It is important to note that we used the same model settings to train both the original and augmented data, ensuring a fair comparison.

For all methods, we preprocessed microbiome features using centered log-ratio transformation (CLR) (Aitchison, 1982) prior to data augmentation. To showcase the broad applicability of PhyloMix, we also conducted experiments using relative abundance data, which was obtained by normalizing the original count data without applying a centered log-ratio transformation. In this setup, we replaced the linear kernel in the SVM with a Radial Basis Function (RBF) kernel to better capture the characteristics of the data.

### A.4 Polytomy resolution

PhyloMix uses the Web of Life (WoL) phylogeny (Zhu et al, 2019), which includes 15, 953 microbial genomes. Note that a small number of internal nodes in the WoL phylogeny have more than two lineages (*i*.*e*., polytomies). The process of resolving polytomies, *i*.*e*., converting them into bifurcating phylogenies, can be either stochastic or deterministic. The stochastic approach arbitrarily assigns a dichotomous structure to nodes with more than two lineages, while the deterministic approach systematically creates a dichotomous structure, beginning with the leftmost child node. As shown in Fig. A.2, the performance of PhyloMix remains robust regardless of the polytomy resolution strategy applied to the phylogeny.

**Fig. A2:**
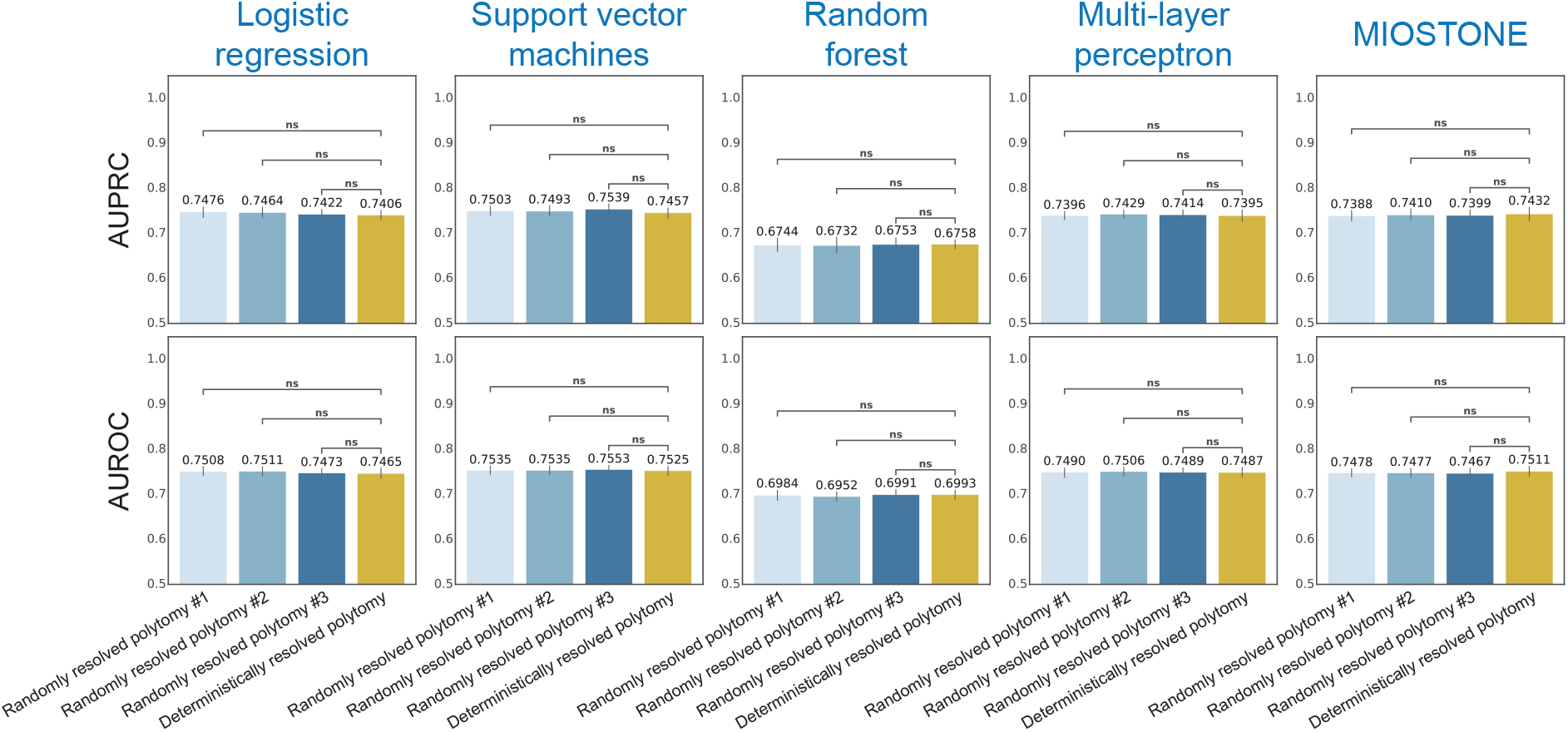
PhyloMix’s performance remains robust regardless of the polytomy resolution strategy applied to the phylogeny. The evaluation was conducted on the IBD dataset and compared against five distinct baseline methods. The WoL phylogeny contains a small number of internal nodes having more than two lineages (*i*.*e*., polytomies), which is resolved using either stochastic or deterministic. To ensure scientific rigor, the stochastic polytomy resolution is performed three times using different random seeds. PhyloMix with different polytomy resolution strategies are measured by both AUPRC and AUROC.

### A.5 Alpha and beta diversity preservation

We calculated the alpha and beta diversity of both the original and PhyloMix-augmented data across six publicly available microbiome datasets. This approach aimed to evaluate whether the augmented data accurately captured the overall correlation structure patterns of the original data. We used the Shannon Index to measure alpha diversity and the Bray-Curtis metric for beta diversity. As shown in Fig. A.3, PhyloMix’s augmented data preserves both the alpha diversity and beta diversity of the original data across six real datasets. We calculated the p-value for alpha diversity using the Kruskal-Wallis test and for beta diversity using PERMANOVA, with the results presented in Tab. A.2.

**Fig. A3:**
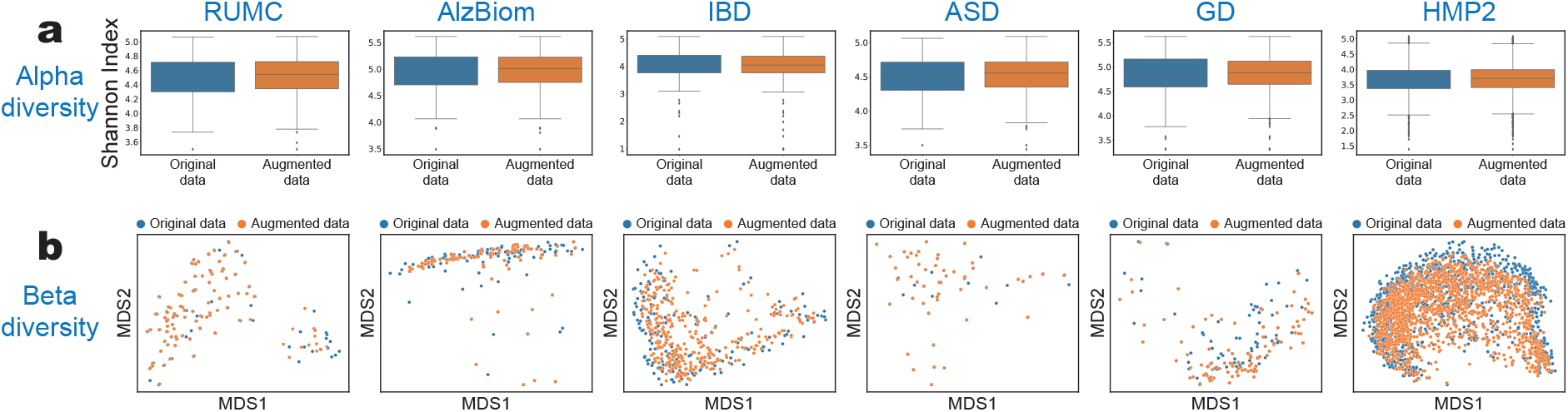
PhyloMix’s augmented data preserves both the alpha diversity and beta diversity of the original data. (**a**) The box plots of the alpha diversity across six real datasets, calculated for each sample in the original data and the augmented data generated by PhyloMix. (**b**) Beta diversity is calculated for every pair of samples from either the original data or the augmented data. The resulting pairwise diversity matrix is visualized in a 2D space using multidimensional scaling (MDS).

**Table A2:** P-values for alpha diversity and beta diversity between the original data and PhyloMix’s augmented data.

| Diversity Metric | IBD | AlzBiom | ASD | RUMC | GD | HMP2 |
| --- | --- | --- | --- | --- | --- | --- |
| Shannon Index | 0.7105 | 0.8126 | 0.9106 | 0.5904 | 0.7288 | 0.2126 |
| Bray-Curtis | 0.9980 | 1.0000 | 1.0000 | 0.9970 | 1.0000 | 1.0000 |

### A.6 Supervised learning results

#### A.6.1 Data with centered log-ratio transformation

**Fig. A4:**
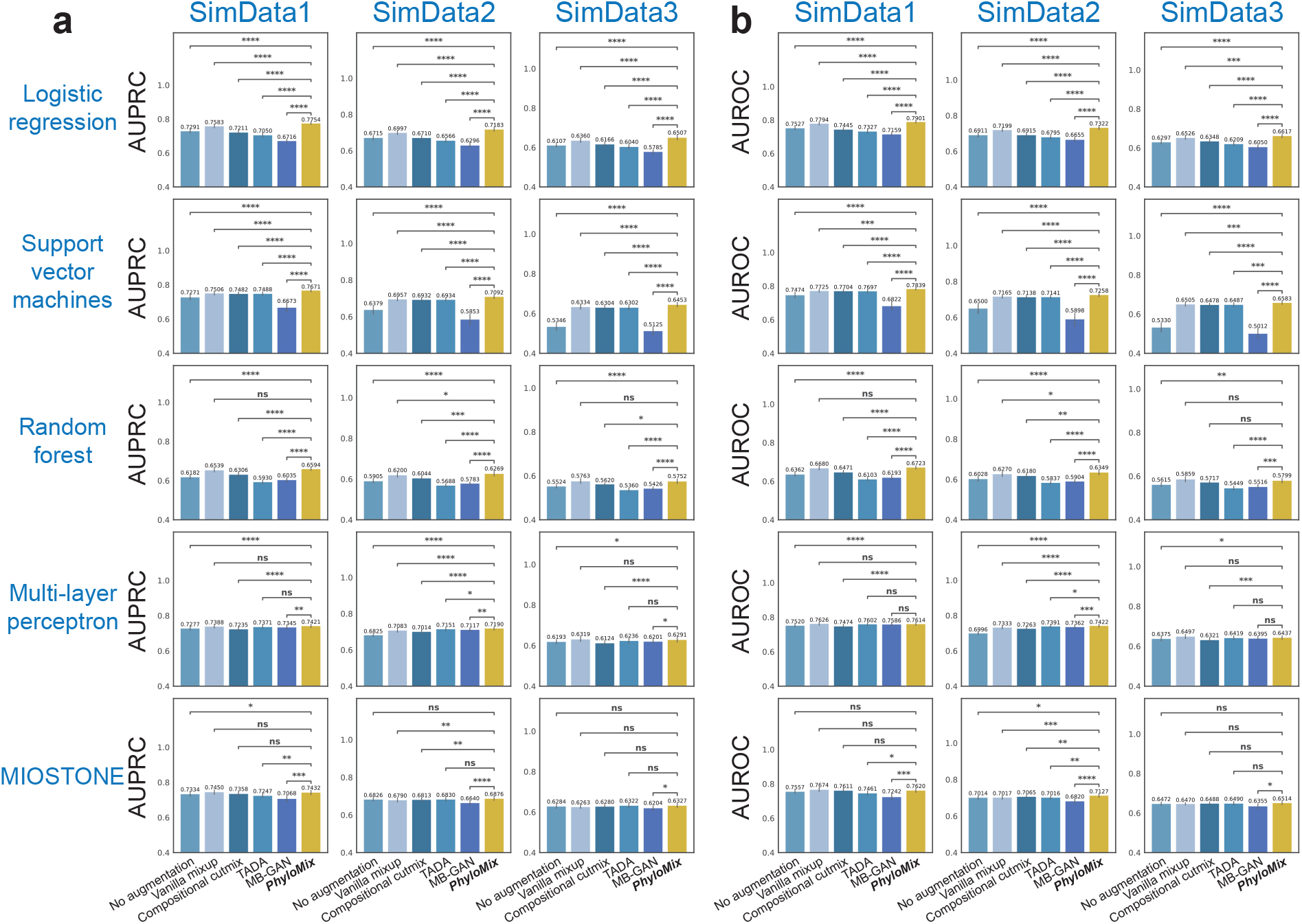
Data augmentation performance on the three simulated datasets in the supervised learning setting. PhyloMix is evaluated alongside five ML models with varying predictive capabilities and compared against four distinct baseline methods. We preprocessed microbiome features using centered log-ratio transformation prior to data augmentation. The performance is measured by (**a**) AUPRC and (**b**) AUROC. For scientific rigor, the performance comparison between PhyloMix and other baseline methods is quantified using one-tailed two-sample t-tests to calculate p-values: ∗ ∗ ∗∗ : p-value ≤ 0.0001; ∗ ∗ ∗ : p-value ≤ 0.001; ∗∗ : p-value ≤ 0.01; ∗ : p-value ≤ 0.05; ns : p-value *>* 0.05.

**Fig. A5:**
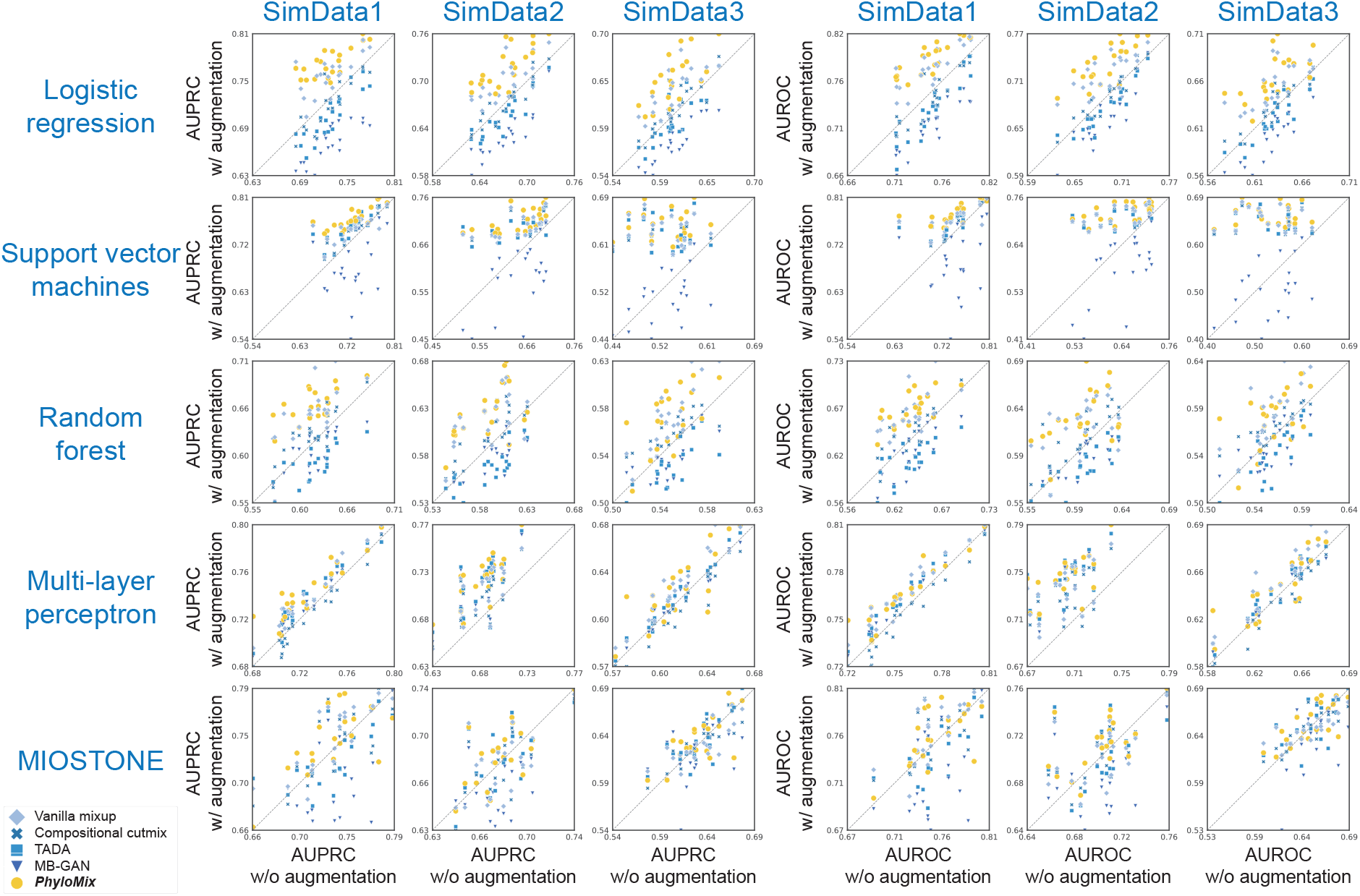
Qualitative evaluation of PhyloMix’s performance on the three simulated datasets in the supervised learning setting. Each dot corresponds to the performance with or without data augmentation for a given method under a random seed. The performance is measured by (**a**) AUPRC and (**b**) AUROC.

**Fig. A6:**
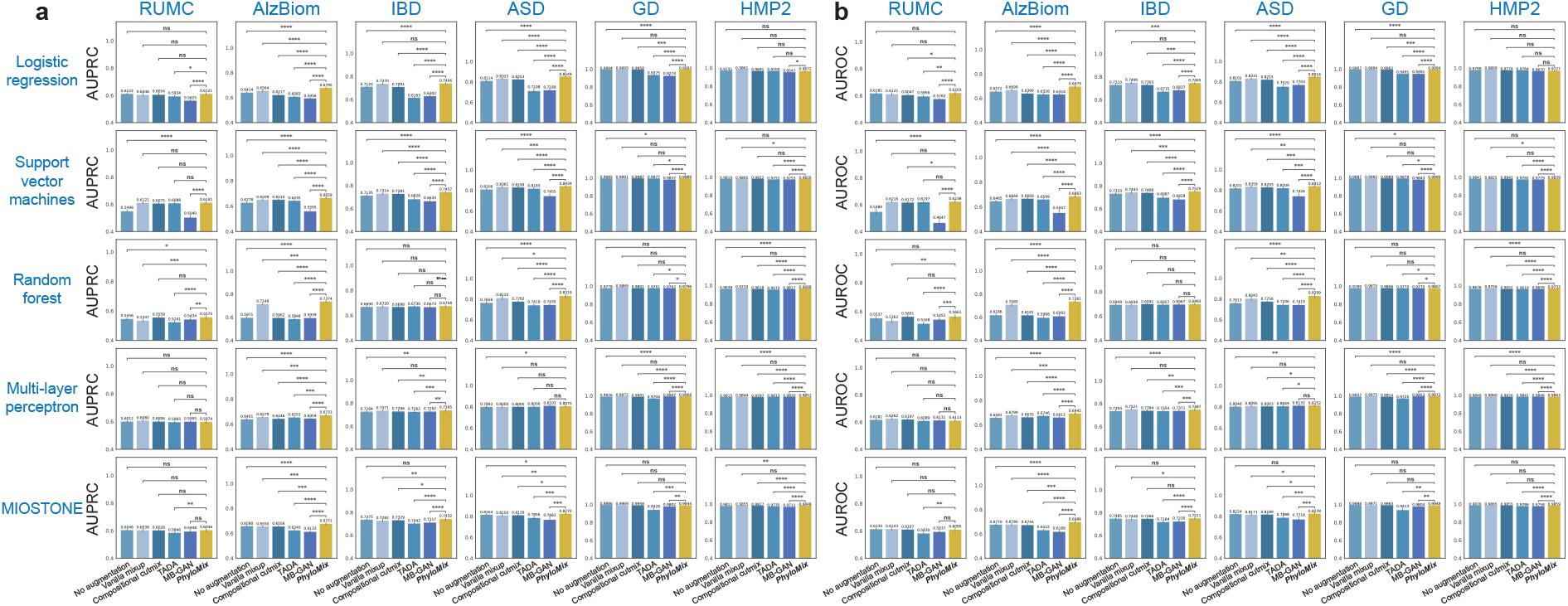
Data augmentation performance on the six real datasets in the supervised learning setting. PhyloMix is evaluated alongside five ML models with varying predictive capabilities and compared against four distinct baseline methods. We preprocessed microbiome features using centered log-ratio transformation prior to data augmentation. The performance is measured by (**a**) AUPRC and (**b**) AUROC. For scientific rigor, the performance comparison between PhyloMix and other baseline methods is quantified using one-tailed two-sample t-tests to calculate p-values: ∗ ∗ ∗∗ : p-value ≤ 0.0001; ∗ ∗ ∗ : p-value ≤ 0.001; ∗∗ : p-value ≤ 0.01; ∗ : p-value ≤ 0.05; ns : p-value *>* 0.05.

**Fig. A7:**
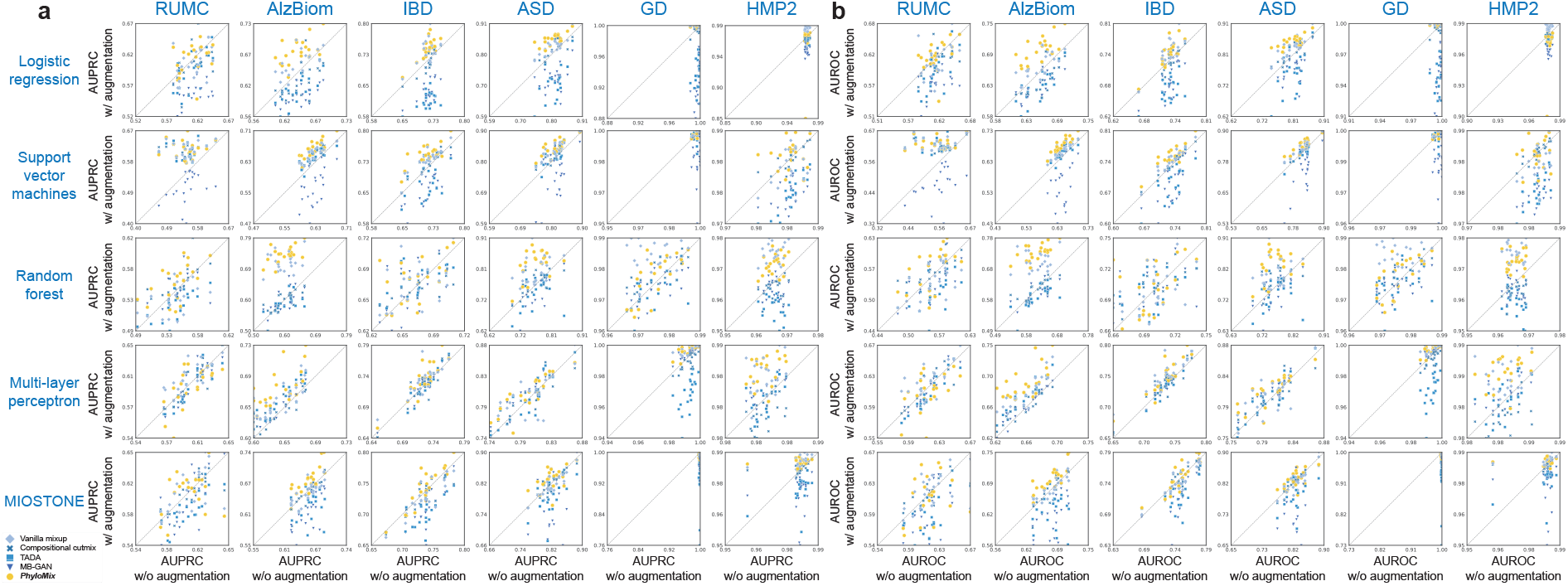
Qualitative evaluation of PhyloMix’s performance on the six real datasets in the supervised learning setting. Each dot corresponds to the performance with or without data augmentation for a given method under a random seed. The performance is measured by (**a**) AUPRC and (**b**) AUROC.

**Fig. A8:**
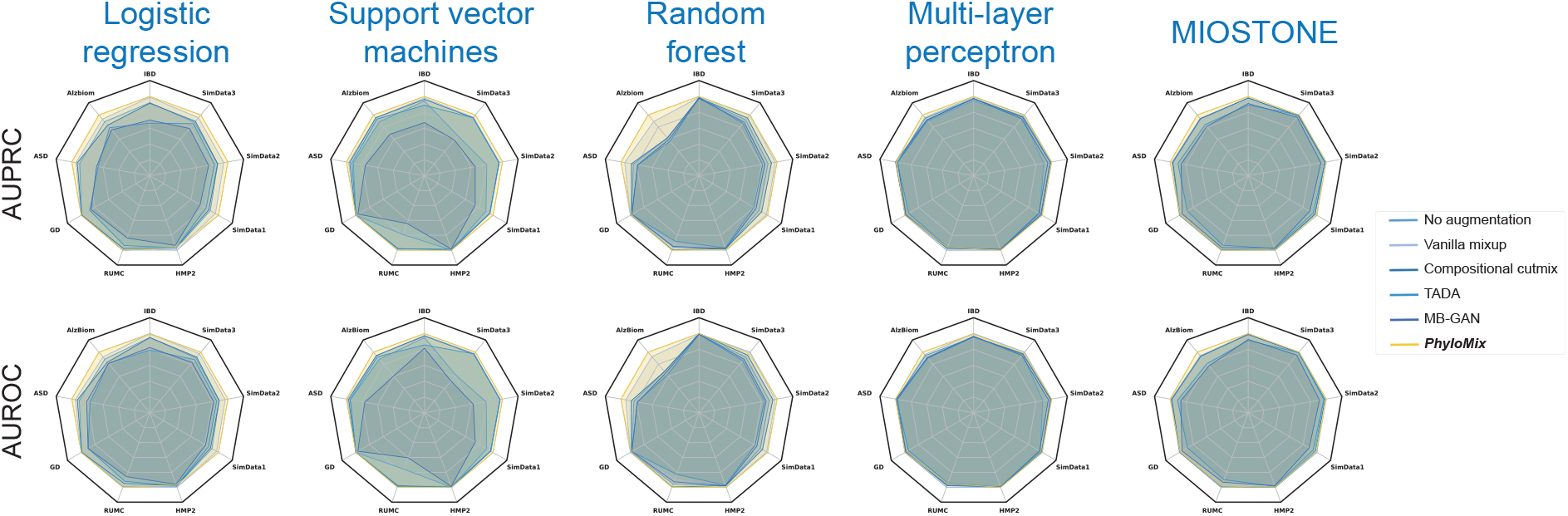
Comparing PhyloMix against baselines methods on all datasets in the supervised learning setting using radar plots. Performance is evaluated using relative AUPRC or AUROC, calculated by normalizing the values against the best performer across all methods.

#### A.6.2 Data with relative abundance normalization

**Fig. A9:**
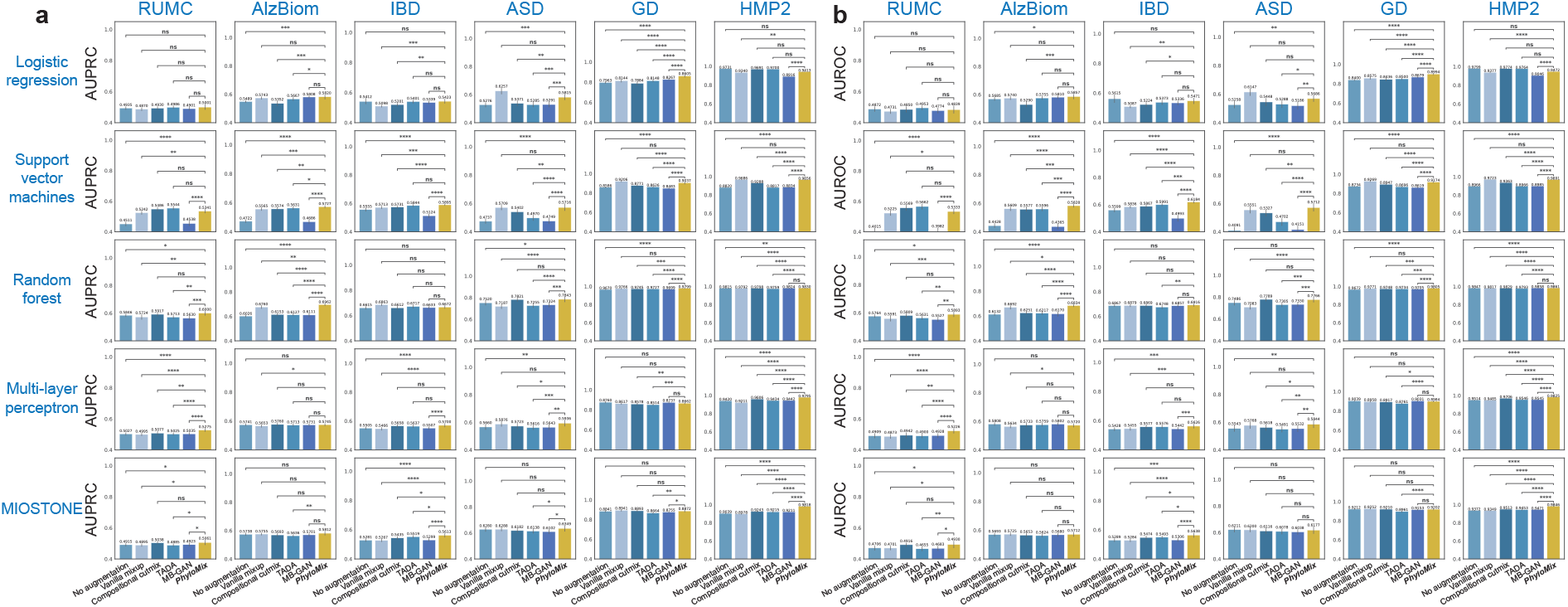
Data augmentation performance on the six real datasets in the supervised learning setting. PhyloMix is evaluated alongside five ML models with varying predictive capabilities and compared against four distinct baseline methods. We preprocessed microbiome features using relative abundance normalization prior to data augmentation. The performance is measured by (**a**) AUPRC and (**b**) AUROC. For scientific rigor, the performance comparison between PhyloMix and other baseline methods is quantified using one-tailed two-sample t-tests to calculate p-values: ∗ ∗ ∗∗ : p-value ≤ 0.0001; ∗ ∗ ∗ : p-value ≤ 0.001; ∗∗ : p-value ≤ 0.01; ∗ : p-value ≤ 0.05; ns : p-value *>* 0.05.

### A.7 Ablation results

**Fig. A10:**
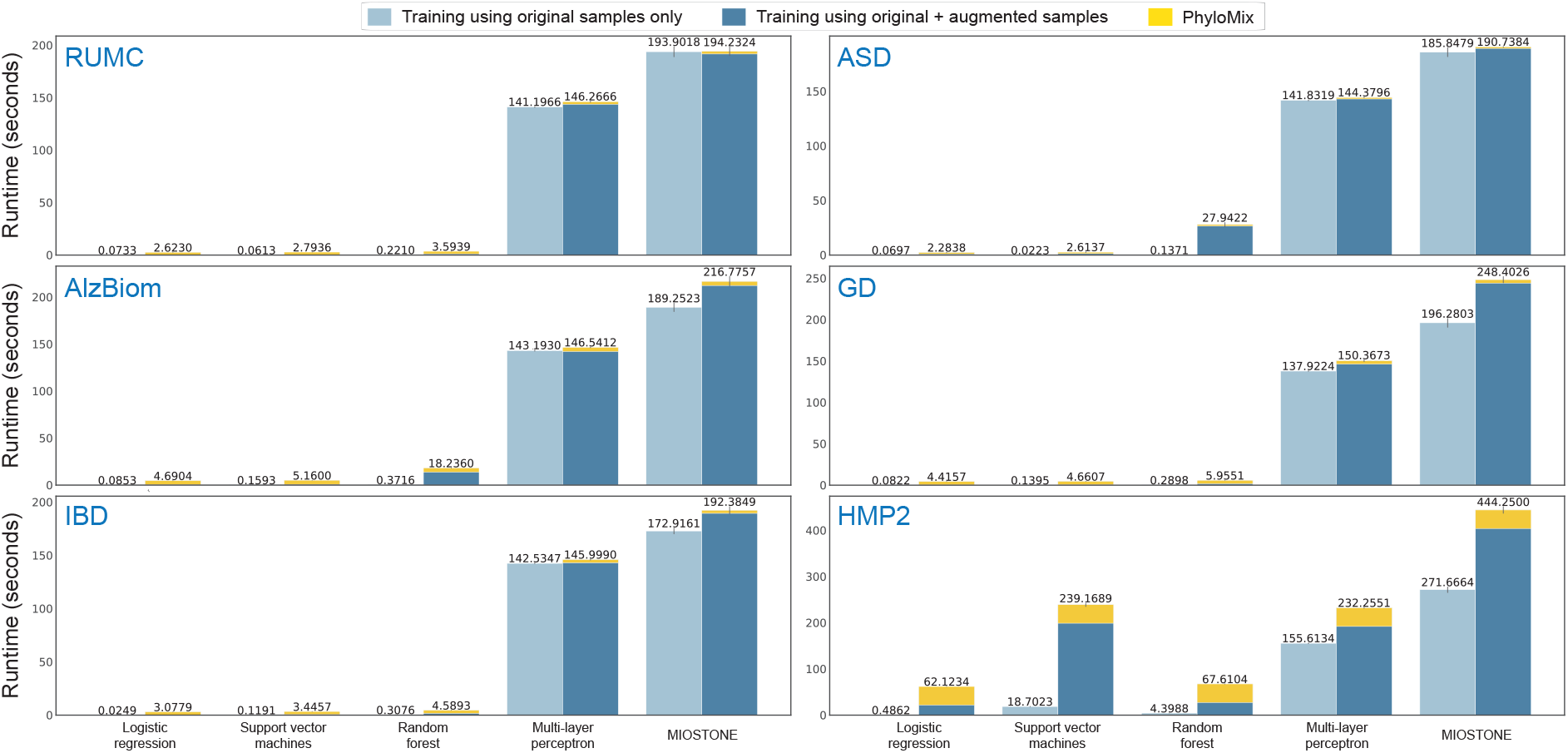
The computational cost of PhyloMix.

**Fig. A11:**
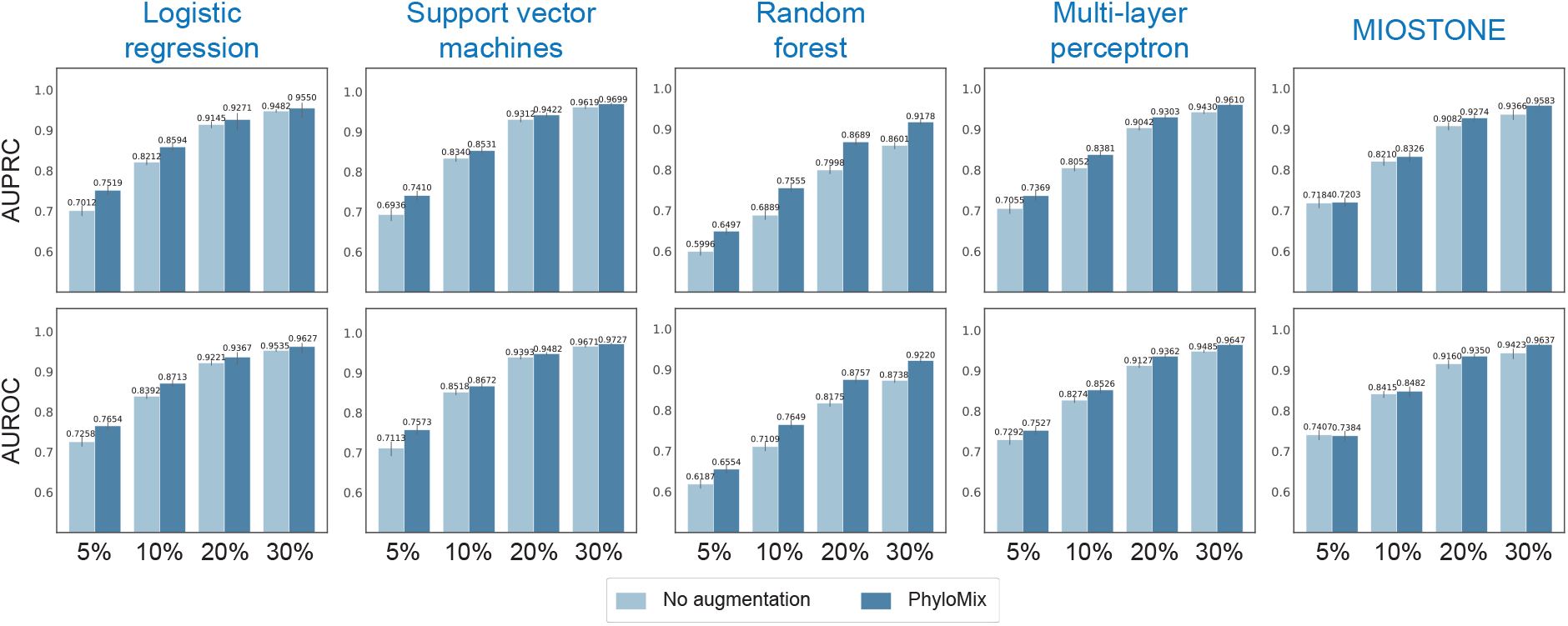
PhyloMix exhibits more significant improvements when the training data size is small.

